# Cyanogenic millipede genome illuminates convergent evolution of cyanogenesis-related enzymes

**DOI:** 10.1101/2025.03.24.645106

**Authors:** Takuya Yamaguchi, Yasuhisa Asano

## Abstract

Hydrogen cyanide (HCN) is a highly toxic biogenic compound. Unlike most natural defensive chemicals, which are typically lineage-specific, the biosynthesis and liberation of HCN, called “cyanogenesis”, occur sporadically among arthropod and plant lineages. This suggests that cyanogenesis has evolved independently numerous times in the animal and plant kingdoms. Although cyanogenesis was identified in millipedes 140 years ago, the cyanogenesis-related enzymes in these arthropods remain unknown. Here, we report a complete set of cyanogenesis-related enzymes in the millipede *Chamberlinius hualienensis* based on an analysis combining genome sequencing and biological characterisation. The gene encoding hydroxynitrile lyase, which catalyses the liberation of HCN from (*R*)-mandelonitrile, and its paralogous genes were clustered, indicating sequential duplication of their coding genes, giving rise to hydroxynitrile lyase in millipedes. We discovered that (*R*)-mandelonitrile biosynthesis in *C. hualienensis* utilises a flavin-dependent monooxygenase (ChuaMOxS) for the initial aldoxime synthesis step, similar to the process in ferns, instead of cytochrome P450 (CYP) as in higher plants and insects. Furthermore, although a single CYP is responsible for converting aldoxime into cyanohydrin in plants and insects, the reaction involves two enzymes in millipedes. We found two CYPs (CYP4GL4 and CYP30008A2) that catalyse aldoxime dehydration to produce nitrile, in addition to CYP3201B1, which catalyses the formation of (*R*)-mandelonitrile from nitrile. The discovery of cyanogenesis-related enzymes in millipedes demonstrates that cyanogenic millipedes evolved these enzymes independently from plants and insects.

**Significance Statement:** The biosynthesis of natural defensive chemicals is usually lineage-specific; however, cyanogenesis (hydrogen cyanide biosynthesis) occurs sporadically among animal and plant lineages. This suggests that the cyanogenesis pathway has arisen numerous times in different kingdoms; however, examples of the independent evolution of the entire pathway are rare. Based on genome sequencing analysis, we report a set of cyanogenesis-related enzymes in the millipede *Chamberlinius hualienensis*. Our findings demonstrate that cyanogenic millipedes evolved independently from plants and insects, providing a deeper understanding of the mechanisms underlying the evolution of metabolic pathways.

## Introduction

Chemical substances, so-called secondary or specialised metabolites, are involved in the most important biotic interactions between plants and their herbivores/pathogens and between animals and their predators/parasites (1). Selection for increased fitness has resulted in each living organism synthesising a distinct set of specialised metabolites appropriate for its environment (2). The acquisition of new enzymes with new functions in specialised metabolite biosynthesis is generally a result of divergent evolution. Thus, most specialised metabolites are typically lineage-specific. In contrast, convergent evolution has been reported, in which different lineages independently evolved the ability to synthesise identical specialised metabolites (1, 3, 4). However, it is less common and less well understood than divergent evolution (5).

Many chemical substances produced by plants and animals play crucial roles in defence. Among the most deterring and toxic biogenic substances, hydrogen cyanide (HCN) inhibits mitochondrial cytochrome *c*-oxidase in the cellular respiration system, limiting the organism’s ability to use oxygen (6). The biosynthesis and liberation of HCN, known as “cyanogenesis”, are widespread among plants (7). Cyanogenic plants accumulate cyanogenic glycosides as stable cyanide precursors. When plant tissues are disrupted by herbivores or pathogens, glycosides are degraded by β-glycosidase and hydroxynitrile lyase (HNL) to release aldehydes or ketones and HCN via cyanohydrins.

Cyanogenesis has been observed in several arthropods, including millipedes, mites, beetles, true bugs, and butterflies. Cyanogenesis was first discovered in millipedes in 1882 (8). Among millipedes, Polydesmida are generally cyanogenic (9). These millipedes accumulate (*R*)-mandelonitrile (MAN) as a cyanide precursor in the reservoirs of defensive glands housed in the paratergas and eject cyanogenic secretions through small openings called ozopores on the dorsal surface near the tips of paired notal projections (Fig. 1A). Despite being identified over 100 years ago, to our knowledge, associated enzymes have yet to be characterised in millipedes, likely owing to the relative rarity of millipedes for collection. However, the Polydesmida millipede *Chamberlinius hualienensis* Wang, which invaded Japan from Taiwan, often forms large swarms (10). This is beneficial for research purposes but can cause problems in local communities because these swarms can enter houses and sometimes cause train delays (11).

**Figure 1.**
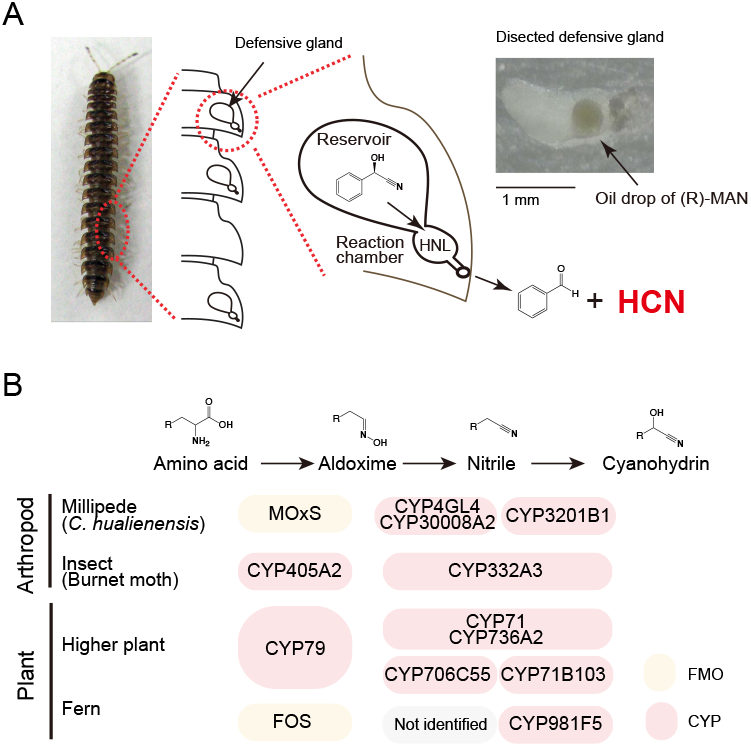
Schematic overview of cyanogenesis in *Chamberlinius hualienensis* and cyanohydrin biosynthesis in millipedes, insects, and plants. (A) *C. hualienensis* and a schematic view of cyanogenesis. Paratergas of the millipede house defensive glands and consist of reservoir and reaction chamber. (*R*)-Mandelonitrile (MAN) stored in the reservoir is admitted to the reaction chamber, converted by hydroxynitrile lyase (HNL) to hydrogen cyanide (HCN) and benzaldehyde, and released to the outside via ozopores. (B) In arthropods (millipede and burnet moth) and plants, cyanohydrin is commonly synthesised from amino acids via aldoxime and nitrile. Different classes of flavin-dependent monooxygenases (FMO) and cytochrome CYPs are responsible for the biosynthesis.

We previously purified HNL (ChuaHNL) from 29 kg of *C. hualienensis* as a model (12). HNL shared no similarity with other proteins but was conserved among cyanogenic millipedes (13, 14). HNL is involved in liberating HCN and benzaldehyde from (*R*)-MAN in millipedes (Fig. 1A). It also catalyses the reverse reaction, namely, the asymmetric synthesis of chiral cyanohydrin, which is a valuable building block for the synthesis of pharmaceuticals from various aldehydes and cyanide sources. ChuaHNL showed the highest specific activity (7420 U/mg) for (*R*)-MAN synthesis among the HNLs isolated from plants and microorganisms. ChuaHNL also exhibited excellent enantioselectivity towards various substrates and high heat and pH stability. Optically pure cyanohydrins are valuable building blocks; therefore, millipede HNLs are used to produce valuable substances (13, 15).

(*R*)-MAN is biosynthesised from L-phenylalanine (Phe) via (*E/Z*)-phenylacetaldoxime (PAOx) and phenylacetonitrile (PAN) in the millipede. The sequence of biosynthesis to cyanohydrin from amino acid via aldoxime and nitrile corresponds to that of cyanohydrin biosynthesis in other cyanogenic plants and arthropods (Fig. 1B). In higher plants, two multifunctional CYPs, CYP79 and CYP71, are generally involved in cyanohydrin biosynthesis with several exceptions (Fig. 1B) (16). The burnet moth (*Zygaena filipendulae*) is the only arthropod for which a complete set of biosynthetic enzymes has been identified (that is, CYP405A2 and CYP332A3; Fig. 1B) (17). Higher plant and insect CYPs catalyse the same reactions but are not homologous, indicating convergent evolution. Compared to plants or burnet moths, *C. hualienensis* may utilise a different set of enzymes. CYP3201B1 from *C. hualienensis* synthesises (*R*)-MAN from PAN but not (*E/Z*)-PAOx (18). Thus, the conversion of aldoxime to cyanohydrin is catalysed by a single CYP in most higher plants and burnet moths but by multiple enzymes in *C. hualienensis*. However, aldoxime- and nitrile-producing enzymes have not yet been identified in millipedes.

Given the scattered distribution of cyanogenesis and the phylogenetic distribution of millipedes, insects, and plants, millipedes independently invented cyanogenesis-related enzymes from insects and plants while the pathway sequence was conserved. In this study, we aimed to identify the enzymes involved in the cyanogenesis pathway in the cyanogenic millipede *C. hualienensis* through genome sequencing analysis. We then biochemically characterised these enzymes.

## Results

### Sequencing and assembly of the *C. hualienensis* genome

In total, 100.42 Gb (∼700× coverage) of clean data from Illumina and 20.04 Gb (∼140× coverage) of clean data from PacBio Sequel were obtained (Tables S1 and S2). A schematic diagram of the hybrid assembly is shown in Fig. S1. The assembled *C. hualienensis* genome was 143,521,810 bp, with a scaffold N50 size of 12,989,427 bp (Fig. 2A and Table 1), representing 99.3% of the estimated genome size (144.5 Mb) (Fig. S2). The integrity of the assembled genome was assessed. First, the short reads obtained from the Illumina sequencing data were compared with the assembled genome. The percentage of reads mapped to the draft genome was >96%. Next, BUSCO (19) analysis revealed that 95.6% of the single-copy arthropod orthologues were complete. These results indicated that the *C. hualienensis* genome assembly was of high quality and coverage. The genome was the smallest among the genome-sequenced millipedes and centipedes, ranging from 181 to 2530 Mbp (Fig. S3).

**Table 1.** Assembly and annotation statistics of *Chamberlinius hualienensis* genome.

| Assembly features | Statistics |
| --- | --- |
| BioProject Accession number | PRJDB13209 |
| Estimate genome size (Mb) | 144 |
| Number of contigs | 104 |
| Contig N50 (Mb) | 4.46 |
| Number of scaffolds | 55 |
| Scaffold N50 (Mb) | 13.0 |
| Assembly size (bp) | 143,521,810 |
| GC content (%) | 31.9 |
| Complete ratio of BUSCO (%) | 95.6 |
| Repeat content (%) | 17.7 |
| Gene number | 14132 |
| Average gene length (bp) | 5971 |
| Average coding sequence length (bp) | 1550 |

**Figure 2.**
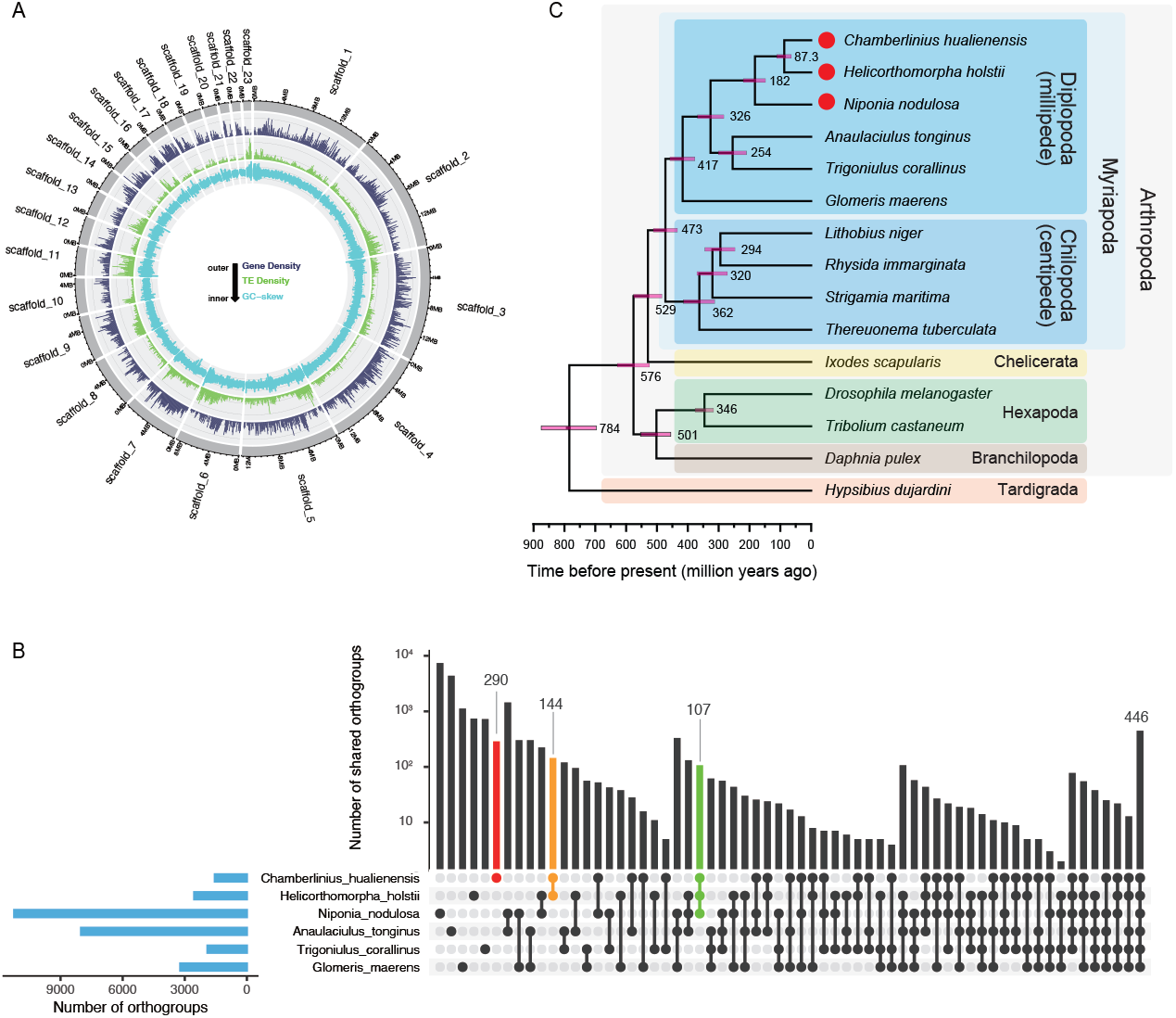
*C. hualienensis* genome annotation. (A) Characterisation of different elements on the millipede scaffolds. From outer to inner ring, the >10 kb scaffolds of the assembled *C. hualienensis* genome, gene density, transposable element density, and GC-skew. (B) Upset plot comparing shared orthogroup in millipedes. (C) Phylogenetic tree. Estimated divergence times (in millions of years) are shown below the phylogenetic tree in black. The tree is rooted with *Hypsibius dujardini* as the outgroup.

For the *C. hualienensis* genome, 14,132 gene models were predicted (Table 1). BUSCO assessment indicated that the predicted genes covered 93.8% of arthropod-conserved orthologues. Among the predicted proteins, 12,443 (80.3%) were annotated using GenBank number, 12,541 (81.2%) using interproscan, 11,702 (75.7%) using eggnog mapper, and 11,038 (71.4%) using pfam.

In the *C. hualienensis* genome assembly, 25.4 Mb of repetitive sequences, which cover 17.7% of the genome, were identified (Table S3). The unclassified elements were the most abundant type of repetitive sequence in *C. hualienensis*, spanning 15.8 Mb (11.0%) of the genome. Class I long terminal repeat retrotransposons account for 1.55% of the genome and comprise the endogenous retrovirus groups ERVL, ERVL-MaLR, ERVclassI, and ERV_classII. Non-long terminal repeat elements account for approximately 0.25% of the genome and comprise short interspersed nuclear elements, ALUs, and the long interspersed nuclear element superfamilies LINE1, LINE2, and L3/CR1 clades. Class II DNA transposons (3.26%) comprised hAT-Charlie and TcMartiggers. The size of the repeat genome sequence, comprising transposable elements and simple repeats, was positively correlated with genome size. Strong correlations between transposable element content and genome size among myriapod genomes were observed (R^2^ = 0.98, *P* = 7.85e-08; regression line y = 128 + 1.63 x) (Fig. S3).

### Orthogroup and species tree

Across myriapod and other arthropod species: millipedes (*Helicorthomorpha holstii, Nipponia nodulosa, Anaulaciulus tonginus, Trigoniulus corallinus*, and *Glomeris maerens*), centipedes (*Rhysida immarginata, Lithobius niger, Thereuonema tuberculate*, and *Strigamia martima*), other arthropods (*Ixodes scapularis, Daphnia pulex, Tribolium castaneum*, and *Drosophila melanogaster*), and the tardigrade *Hypsibius dujardini*, 41,810 orthogroups were identified. In total, 3,104 orthogroups contained all species, 18,160 contained a subset of species, and 21,323 were species-specific orthogroups. Six millipede species shared 446 orthogroups, three Polydesmida millipede species shared 107 orthogroups, two cyanogenic millipedes shared 144 orthogroups, and *C. hualienensis* shared 290 species-specific orthogroups (Fig. 2B). To infer the evolutionary history of *C. hualienensis*, phylogenetic reconstruction and divergence time estimation analyses were conducted based on 102 single-copy orthologous proteins shared by the abovementioned organisms (Fig. 2C). The estimated divergence time for Diplopoda was approximately 473 Mya (Middle Ordovician period), with a 95% highest posterior density (HPD) of 433–512 Mya, which aligns with the previously estimated divergence time (468–518 Mya) (20). Three Polidesmida species, *C. hualienensis, H. hostii*, and *N. nodulosa*, formed a monophyletic clade with an estimated divergence time of 326 Mya (Carboniferous period) and a 95% HPD of 282–369 Mya. The divergence time between *C. hualienensis* and *H. holstii* was approximately 87.3 Mya (Late Cretaceous period), with a 95% HPD of 64.1–111 Mya.

### *ChuaHNL* and its paralogous gene cluster on the genome

The millipede accumulates (*R*)-MAN (males 564 ± 269 and females 596 ± 137 µg/millipede) as a cyanide precursor (Fig. S4), and ChuaHNL catalyses the release of HCN from the compound (21). *ChuaHNL* (CHUA_005138) and its two paralogues (CHUA_005137 and CHUA_005136) were grouped into the orthogroup OG0016952. These protein-coding genes were clustered in the genome (Fig. 3-A). CHUA_005137 and CHUA_005136 shared 70% and 54% amino acid sequence identities with ChuaHNL, respectively (Fig. S5), and shared all and four of the five catalytic residues, respectively (Fig. 3-B and Fig. S6). These results suggested that ChuaHNL paralogous proteins exhibit HNL activity.

**Figure 3.**
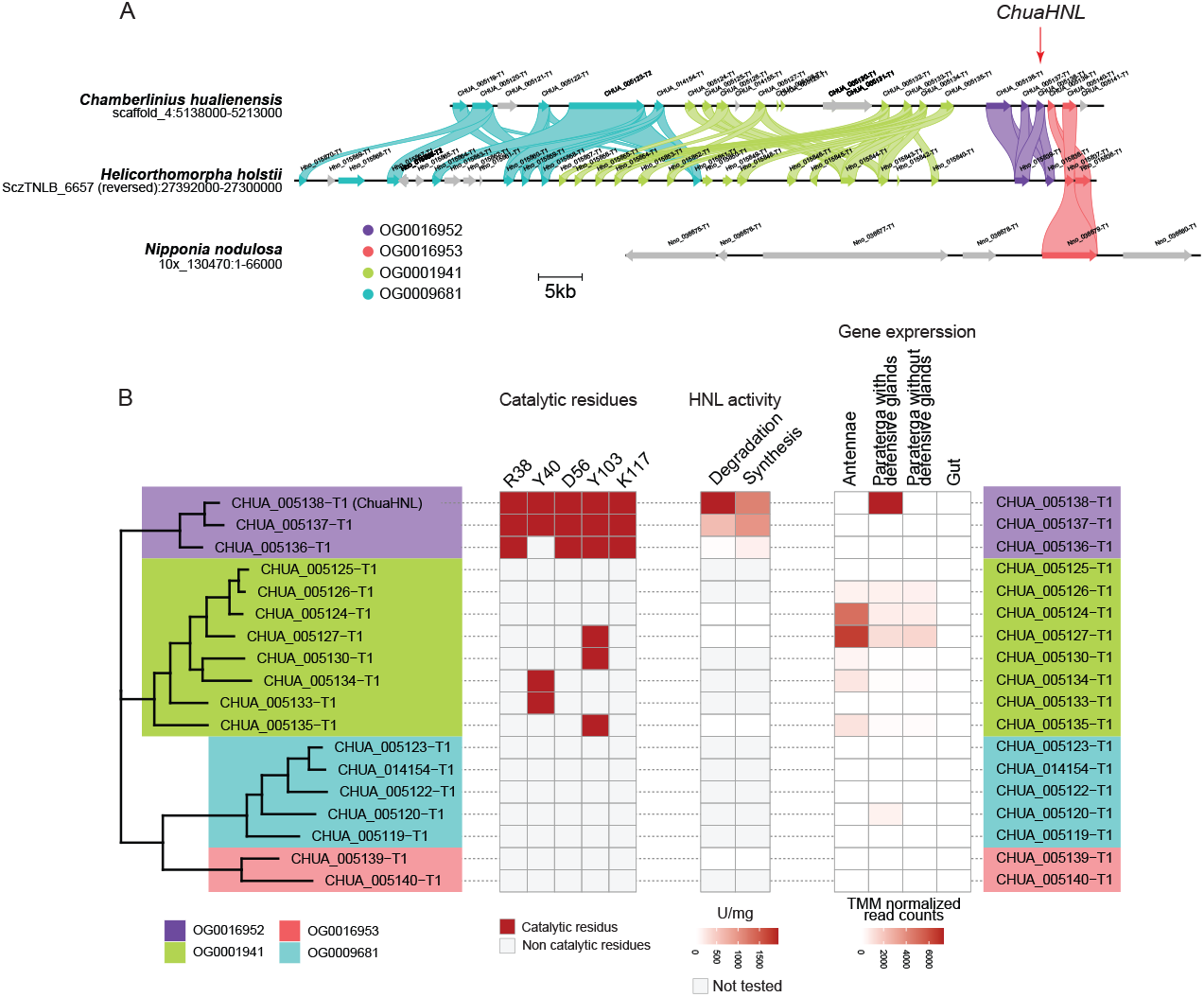
Gene cluster and characterisation of *ChuaHNL* and its paralogues. (A) *ChuaHNL* and its paralogue cluster on the *C. hualienensis* genome and synteny among Polydesmida millipedes. (B) Phylogenetic tree and catalytic residues, HNL activity, and gene expression (TMM-normalised reads counts) of *ChuaHNL* and its paralogues.

For biochemical characterisation, we heterologously produced ChuaHNL, CHUA_005137, and CHUA_005136 as N-terminal His-tagged proteins using the yeast *Pichia pastoris* expression system (22). ChuaHNL, CHUA_005137, and CHUA_005136 were isolated from 120 mL of the culture supernatant using a nickel affinity column. SDS-PAGE showed that all proteins were secreted into the medium (Fig. S7). These three proteins catalysed (*R*,*S*)-MAN degradation, indicating that CHUA_005137 and CHUA_005136 are functional HNLs. To characterise the paralogous proteins, recombinant proteins were purified from 1.5 L of culture supernatant (Tables S4–S6). ChuaHNL and two paralogous proteins catalysed the degradation of racemic MAN into benzaldehyde and HCN with the following kinetic parameters (Fig. S8-A): ChuaHNL, *V*_max_ = 2950 ± 227 µmol/min/mg, *K*_m_ = 13.7 ± 2.35 mM; CHUA_005137, *V*_max_ = 916 ± 67.4 µmol/min/mg, *K*_m_ = 12.9 ± 2.16 mM; CHUA_005136 *V*_max_ = 50.0 ± 0.681 µmol/min/mg, *K*_m_ = 1.10 ± 0.0618 mM. ChuaHNL, CHUA_005137, and CHUA_005136 also catalysed its reverse reaction, namely the (*R*)-MAN synthetic reaction from benzaldehyde and potassium cyanide, with the following kinetic parameters: ChuaHNL, *V*_max_ = 1310 ± 63.6 µmol/min/mg, *K*_m_ = 4.31 ± 0.771 mM; CHUA_005137 *V*_max_ = 1280 ± 80.1 µmol/min/mg, *K*_m_ = 4.50 ± 0.994 mM; CHUA_005136 *V*_max_ = 677 ± 93.6 µmol/min/mg, *K*_m_ = 2.36 ± 0.696 mM, *K*_i_ = 21.1 ± 5.74 mM. When the same activity of the enzyme was used for the reaction (that is, 8 U/mL), the enantiomeric excess of (*R*)-MAN produced was 92.1%, 91.5%, and 83.8%, respectively (Fig. S9). These results indicated that CHUA_005137 and CHUA_005136 have excellent *R*-selectivity in cyanohydrin synthesis, similar to ChuaHNL (12).

Seventeen *ChuaHNL*-like genes were observed within 75 kb of the *ChuaHNL* gene cluster (Fig. 3-A). The proteins encoded by these genes were assigned to orthogroups OG0001941, OG0009681, and OG0016953, unlike ChuaHNL, which belonged to OG0016952 (Fig. 3-A and - B). However, the amino acid sequences of these proteins shared 11–21% identity at the amino acid level with ChuaHNL (Fig. S5) and eight structurally important Cys residues (Fig. S6), which formed inter- and intra-disulfide bonds in ChuaHNL (23), except *CHUA_005132*. This gene was partially truncated and is considered malfunctional (Fig. S6). These results suggested that ChuaHNL and the paralogous proteins, except for CHUA_005132, share similar three-dimensional structures, although these paralogous proteins are grouped into four orthogroups. We evaluated the HNL activity of several paralogous proteins (CHUA_005139 from OG0014028; CHUA_005135, CHUA_005127, and CHUA_005124 from OG000814). These proteins were successfully produced in *P. pastoris* (Fig. S7) but did not show HNL activity (Fig. 3-B), which agrees with the fact that these paralogous proteins did not have the catalytic residues of millipede HNLs (Fig. 3-B and Fig. S6). We attempted to create variants with full active site residues but were unable to do so in *P. pastoris*.

*ChuaHNL* was specifically expressed in the paraterga-containing defensive glands (Fig. 3-B) in accordance with previous gene expression analyses (12). CHUA_005137 and CHUA_005136 were expressed in traces, indicating that these proteins do not accumulate in the defence glands, although they exhibit HNL activity. A few paralogous proteins, such as CHUA_005124 and CHUA_005127, were highly expressed in the antennae (Fig. 3-B). ChuaHNL belongs to the lipocalin family and functions as a carrier protein for small hydrophobic molecules. The expression of ChuaHNL paralogues in the antennae is consistent with that of lipocalin genes in the antennae of several insect species.

Orthologues of *ChuaHNL* and its paralogous genes were found in the *H. holstii* genome. They clustered in the *C. hualienensis* genome, and their syntheny was conserved (Fig. 3-A). The *N. nodulosa* genome only encoded an orthologous protein belonging to OG001428, not the gene cluster in *C. hualienensis* and *H. holstii* (Fig. 3-A).

### CYPome and identification of CYPs involved in (*R*)-MAN biosynthesis

CYPs are key biosynthetic enzymes of cyanohydrins in cyanogenic plants and burnet moths (Fig. 1-A) (10, 24). However, the CYPs in Myriapoda are not well understood. We searched for CYPs in millipede and centipede genomes and found 96 *C. hualienensis* genes that encode putative functional CYPs (Fig. 4-A). The Polydesmida millipedes *H. holstii* and *N. nodulosa* contain 119 and 80 CYPs, respectively. These numbers were greater than those of other millipedes (32–64) and centipedes (52–63) (Fig. S10). Phylogenetic analysis revealed that *C. hualienensis* CYPs (ChuaCYPs) clustered into the CYP2, CYP3, CYP4, CYP20, and mitochondrial clans (Fig. 4-A). CYP2 clan members comprised the largest number of CYPs in *C. hualienensis* (Fig. S10). Similarly, CYP2 clan members were most abundant in other Myriapoda species (Fig. S10). However, the CYP3 and CYP4 clans are predominant in insects (25). The difference in clan proportions in arthropods is thought to be caused by the dynamics of gene births and deaths over evolutionary time, affecting the CYPome (25).

**Figure 4.**
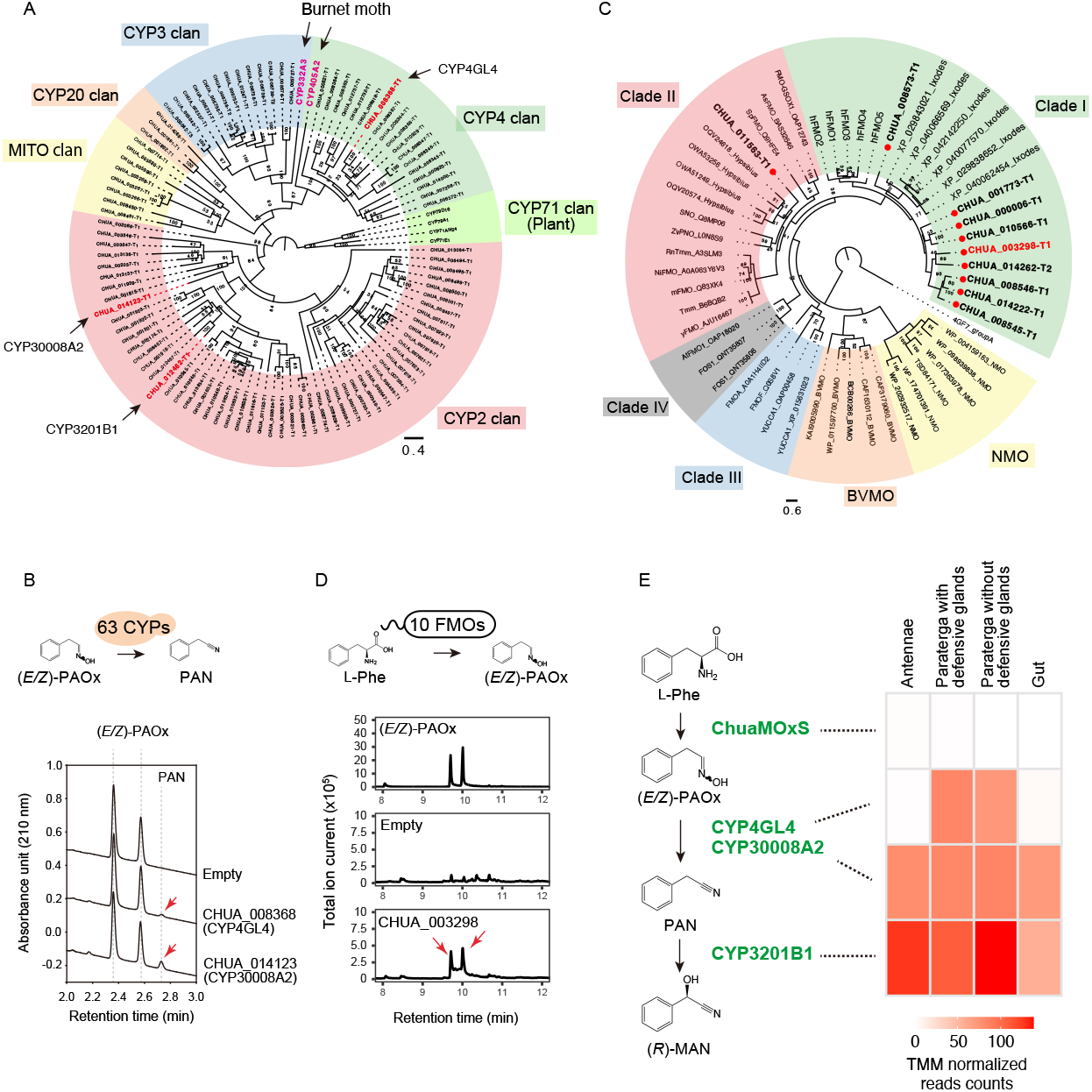
Identification of (*R*)-mandelonitrile biosynthetic enzymes from *C. hualienensis*. (A) Phylogenetic tree of cytochrome P450s (CYPs) from *C. hualienensis* and previously characterised cyanohydrin biosynthetic CYPs from the burnet moth and plants. Bar indicates 40% divergence. (B) Identification of CYPs catalysing the conversion of (*E/Z*)-PAOx into PAN. Yeast cells carrying CYPs from *C. hualienensis* expression plasmids or an empty vector were incubated with (*E/Z*)-PAOx. PAN formation was analysed using ultra-performance liquid chromatography. Reaction product peaks are indicated by red arrows. (C) Phylogenetic tree of FMOs from *C. hualienensis*, arthropods, plants, and microorganisms. Bar indicates 60% divergence. (D) Identification of the (*E/Z*)-PAOx-producing FMOs. *Escherichia coli* cells carrying FMOs from *C. hualienensis* expression plasmids or an empty vector were cultured. The accumulation of (*E/Z*)-PAOx was analysed using gas chromatography–mass spectrometry. Reaction product peaks are indicated by red arrows. (E) (*R*)-Mandelonitrile biosynthetic pathway in *C. hualienensis* and gene expression (TMM-normalised read counts) of biosynthetic enzyme genes.

To narrow down the CYP candidates involved in (*R*)-MAN biosynthesis, we searched for CYPs specifically expressed in paratergas with defensive glands, which accumulate (*R*)-MAN, by analysing expression using RNA-seq. However, we were unable to identify such candidate CYPs. Therefore, we performed functional CYPome analysis to identify (*R*)-MAN biosynthetic CYPs. Of these 88 microsomal CYPs, 63 were cloned. To evaluate the activity of CYPs in yeast, the redox partner cytochrome P450 reductase (CPR) must be co-expressed. In *C. hualienensis*, there is one CPR gene (ChuaCPR, CHUA_001042). We generated a yeast strain, *S. cerevisiae* ChCR11, in which *ChuaCPR* was genomically introduced. The strain exhibited a higher PAN-metabolising activity than the parent strain INVSc1 when expressed *CYP3201B1* (Fig. S11), which catalyses the hydroxylation of PAN to (*R*)-MAN.

We evaluated the activity of 63 ChuaCYPs towards L-Phe and (*E/Z*)-PAOx using whole-cell biocatalysis, which can detect the catalytic activities of CYPs despite their production levels in yeast being below the detection limit of the CO difference-spectrum assay (26). Although no (*E/Z*)-PAOx-producing CYPs were detected, two CYPs, CHUA_008368 and CHUA_014123, produced PAN from (*E/Z*)-PAOx (Fig. 2-B). CHUA_008368 and CHUA_014123 were designated CYP4GL4 and CYP30008A2, respectively, based on the standard CYP nomenclature. CYPs were assigned to the same family and subfamily when the amino acid level sequence identity was >40% and >55%, respectively (27).

Two CYPs, CYP4GL4 and CYP30008A2, belonging to the CYP4 and CYP2 clans, respectively, were phylogenetically distinct. CYP4GL4 showed 65% identity at the amino acid level with CYP4GL3 (CHUA_008364), which did not exhibit nitrile-producing activity. CYP30008A2 showed 74% and 37% identity to CYP30008A1 from the millipede *Polydesmus complanatus* and CYP3201F1 (CHUA_011122), respectively. CYP30008A1 was derived from a transcriptome shotgun assembly sequence archive (GESI01022006.1), and the enzyme was not biochemically characterised. Microsomes harbouring CYP4GL4 and CYP30008A2 catalysed the formation of PAN from (*E/Z*)-PAOx (Fig. 4-B). CYP4GL4 and CYP30008A2 exhibited optimum pH and temperature of 8.0 and 35°C and 7.0 and 40°C, respectively (Fig. S13). The *V*_max_ and *K*_m_ towards (*E/Z*)-PAOx were as follows: CYP4GL4, *V*_max_ = 615 ± 11.6 pmol min^−1^ mg^−1^ (microsome), *K*_m_ = 8.13 ± 1.06 µM; CYP30008A2, *V*_max_ = 106 ± 4.54 pmol min^−1^ mg^−1^ (microsome), *K*_m_ = 164 ± 29.5 µM. Both CYPs act on 4-hydroxyphenylacetaldoxime (4HPAOx) and indole-3-acetldoxime (IAOx) to produce the corresponding nitriles (Fig. S14), indicating that the two nitrile-synthesising CYPs act on multiple amino acid-derived aldoximes.

### Flavin-dependent monooxygenases catalysing the first (*R*)-MAN biosynthesis step

Next, we identified 10 putative functional flavin-dependent monooxygenases (FMOs) in the *C. hualienensis* genome (Fig. 4-C and Table S7) as the candidates for PAOx-producing enzymes. These FMOs had the following FMO signature sequences: FAD-binding motif (GXGXXG), FMO motif (FXGXXXHXXXYK), and NADPH-binding motif (GXGXXG) (Fig. S15-A), and they shared 23– 63% sequence identity (Fig. S15-B). These FMOs, except for CHUA_011563, were predicted to have a transmembrane region at their C-terminus (Table S7) and were localised to the endoplasmic reticulum (Table S7), similar to mammalian FMOs involved in xenobiotic metabolism (28). However, the insect FMOs that have been characterised are soluble proteins (29–31). FMOs were divided into six groups (A–H) based on their structures and properties (32), and group B FMOs were further divided into four clades (33). Phylogenetic analyses showed that the *C. hualienensis* FMOs belonged to clade I in group B, except for CHUA_011563, which belonged to clade II in group B (Fig. 4-D). The other millipede and centipede genomes contained 2–14 genes encoding putative functional FMOs (Fig. S16). These FMOs clustered into clade I or II of group B, as in the case of *C. hualienensis* FMOs (Fig. S16) and were not phylogenetically related to fern FOS1 (clade IV) or microbial FMOs.

All *C. hualienensis* FMOs were expressed and not in a tissue-specific manner. Therefore, all cDNAs encoding FMOs were heterologously produced in *Escherichia coli* to determine the (*E/Z*)-PAOx-producing activity of the FMOs. Among them, *E. coli* harbouring CHUA_003298 accumulated 128 ± 51 µM (*E/Z*)-PAOx in the culture (Figs. 4-D and S17), whereas *E. coli* harbouring the other FMOs did not (Fig. S17-A). Given that CHUA_003298 did not show enzymatic activity after the disruption of *E. coli* cells, biochemical characterisation of the enzyme was not performed. The accumulation of 4HPAOx and IAOx derived from L-tyrosine and L-tryptophan, respectively, was not detected (Fig. S18), indicating that CHUA_003298 has a narrow substrate specificity for producing L-Phe-derived aldoxime. We designated the PAOx-producing enzyme CHUA_003298 as “millipede aldoxime synthase (MOxS)” (Fig. 4-E).

### Distribution of MAN biosynthetic enzyme genes among Myriapod species

Genes encoding cyanogenesis-related enzymes were identified in two Polydesmida millipedes, *H. holstii* and *N. nodulosa*, but not in genome-sequenced non-cyanogenic millipede species from Julida, Spirobolida, and Glomerida nor centipedes (Fig. 5). ChuaMOxS (CHUA_003298) shared 82.2% and 70.6% identity towards Hho_019011 from *H. holstii* and Nno_0338811 from *N. nodulosa*, respectively (Fig. S19). *H. holstii* had CYP4GL4 (Hho_020879) and CYP30008A2 (Hho_011653) orthologues, but the CYP30008A2 orthologue lacked the N- and C-terminal region coding sequences (Fig. S20). Thus, its coding protein is likely malfunctional. CYP4GL4 and Hho_020879 shared 67% identity (Fig. S21). However, *N. nodulosa* harboured the CYP30008A2 orthologue (Nno_033394) but not the CYP4GL4 orthologue. CYP30008A2 and Nno_033394 shared 72% identity (Fig S20). Thus, the nitrile synthesis step likely differs between millipede species that utilise either one or both PAN-producing CYPs (Fig. 5). *H. holstii* and *N. nodulosa* harbour an orthologue of CYP3201B1 that catalyses the conversion of PAN to (*R*)-MAN: Hho_006485 (86%) and Nno_043294 (62%), respectively. These (*R*)-MAN biosynthetic enzymes share relatively high amino acid sequence identity (>65%) among *C. hualienensis, H. holsti*, and *N. nodulosa* but not cyanogenesis-related enzymes from other cyanogenic plants or burnet moths. This indicates that Polydesmida millipedes independently evolved cyanogenesis-related enzyme genes at least 182 Mya (Fig. 2-C).

**Figure 5.**
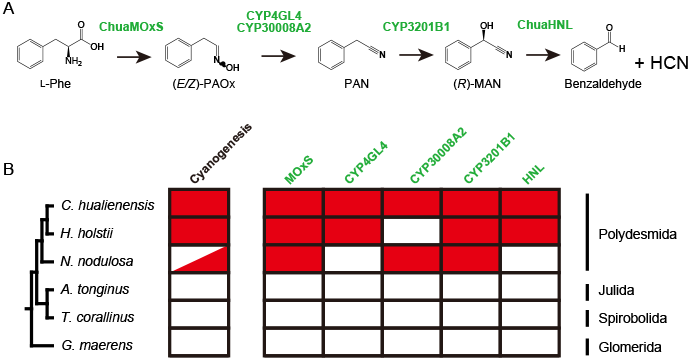
Distribution of cyanogenesis-related enzymes in millipedes. (A) Cyanogenesis pathway in *C. hualienensis*. (B) Red box indicates the presence of orthologues of *C. hualienensis* cyanogenesis-related enzyme genes.

## Discussion

Here, we report on the cyanogenesis pathway of a cyanogenic millipede, *C. hualienensis*. By combining genome sequencing and biochemical characterisation, we identified the enzymes presumably involved in (*R*)-MAN biosynthesis. Given that a set of cyanogenesis-related enzymes are phylogenetically distant, the cyanogenesis pathway likely evolved multiple times across arthropod and plant lineages.

HNL is a key enzyme that releases HCN into cyanogenic millipedes. Millipede HNLs do not exhibit sequence similarity to other proteins (12) but are conserved among cyanogenic millipedes (14). *ChuaHNL* (CHUA_005138) and its paralogous protein-coding genes were clustered (Fig. 3-A). CHUA_005137 and CHUA_005136 exhibited HNL activity similar to those of ChuaHNL. ChuaHNL was highly expressed in the paraterga-containing defensive glands, whereas the expression of these genes was low (Fig. 3-B). Thus, these paralogous proteins are unlikely to accumulate in the reaction chambers of defensive glands. Although the two ChuaHNL paralogous proteins are unlikely to play important roles in millipedes, they catalyse industrially important asymmetric cyanohydrin synthesis with high enantioselectivity (Fig. S9). In particular, the kinetic parameters of CHUA_005137 were comparable with those of ChuaHNL (Fig. S8-B). Millipede HNLs commonly exhibit high specific activity towards (*R*)-MAN synthesis (14), whereas other cyanohydrin synthetic activities differ in terms of specific activity and stereoselectivity (13, 15).

Thus, these ChuaHNL paralogous proteins may have distinct substrate specificities and could be valuable for synthesising optically active cyanohydrins, such as pharmaceutical intermediates. Genomic analysis of cyanogenic millipedes could lead to the discovery of HNLs that are unobtainable by enzyme purification or degenerate PCR-based cloning, as described previously (12–14).

Based on their three-dimensional structure, millipede HNLs belong to the lipocalin family (13, 15, 23). Lipocalins are distributed in all kingdoms of life, except for Archeae (34) and represent a large superfamily of proteins sharing a conserved scaffold; however, amino acid sequences can be highly divergent, with identities as low as 10%. Lipocalins interact with and transport small hydrophobic molecules, such as steroid hormones, odourants (e.g. pheromones), retinoids, and lipids (35). In arthropods, some lipocalin genes are highly expressed in the antennae and pheromone glands and may function as semiochemical carrier proteins (36). Thus, ChuaHNL paralogous proteins may interact with these small hydrophobic molecules. Several paralogues, such as CHUA_005124 and CHUA_005127, are expressed in the antennae (Fig. 3-B), suggesting that they function as chemosensory proteins.

Multiple lipocalin genes were found in arthropod genomes and were clustered on some arthropod species genomes (36, 37), as in the case of the HNL paralogue cluster in *C. hualienensis* and *H. holstii* (Fig. 3-A). The relatively large number of these genes in some arthropod species may have resulted from extensive duplication and differentiation under environmental pressure. Ancestral millipede lipocalin proteins that can bind small hydrophobic molecules may have acquired the catalytic residues necessary for HNL activity through sequential duplication of their coding genes in the genome, giving rise to highly active and stable millipede HNLs.

The pathway for cyanohydrin biosynthesis from amino acids via aldoxime and nitrile is conserved among millipedes, insects, and plants (Fig. 1-B). Class B, clade I FMO ChuaMOxS catalyses the formation of (*E/Z*)-PAOx (Fig. 4-F), the first step in the millipede cyanogenesis pathway. Aldoxime formation in cyanogenic seed plants is generally catalysed by CYPs belonging to the CYP79 family (38). This enzyme catalyses two successive *N*-hydroxylations of the primary amino acid and a decarboxylation step to produce aldoxime. Similarly, CYP405A2 is responsible for the formation of aldoxime in the burnet moth (17). This is an example of convergent evolution across the plant and animal kingdoms. However, some FMOs can produce aldoxime from amino acids, but this has rarely been reported. A group B, clade IV FMO, named FOS1, from a fern, and OOS, from Darwin’s orchid, catalyse the formation of aldoxime from amino acids (39, 40). The microbial FMO SCO7468 from *Streptomyces coelicolor* A3(2) catalyses the conversion of 5-dimethylallyl-L-tryptophan into the corresponding aldoxime (41). As these FMOs are thought to catalyse two successive *N*-hydroxylations, similar to CYP79s, ChuaMOxS presumably produces (*E/Z*)-PAOx via the same reaction mechanism. These millipede, plant, and microbial FMOs are group B FMOs but are phylogenetically distant (Fig. 4-C), indicating that they developed through convergent evolution rather than horizontal gene transfer.

Animal FMOs are xenobiotic-metabolising enzymes with broad substrate specificity that play important roles in drug metabolism, particularly in the human liver (42). Similarly, the insect FMOs characterised thus far catalyse the detoxification of insecticides and toxic plant-specialised metabolites in food (30, 31, 43). In contrast to the previously characterised arthropod FMOs, ChuaMOxS is a narrow substrate-specific enzyme that produces L-Phe-derived (*E/Z*)-PAOx, as it does not produce other amino acid-derived aldoximes (Fig. S17). Furthermore, the expression of ChuaMOxS was considerably lower than that of CYP4GL4, CYP30008A2, and CYP3201B1 (Fig. 4-E). The narrow substrate specificity and gene expression of ChuaMOxS likely indicate the biosynthesis of (*R*)-MAN in millipedes.

Owing to the low level of ChuaMOxS in *E. coli*, detailed functional analysis could not be performed, but other heterologous expression systems such as insect cultured cells or MOxS from other cyanogenic millipedes may help in the detailed characterisation of MOxS.

The second step in (*R*)-MAN biosynthesis is the conversion of (*E/Z*)-PAOx to PAN. We identified two novel CYPs, CYP4GL4 and CYP30008A2, which catalyse the formation of PAN from (*E/Z*)-PAOx (Fig. 3-B). Aldoxime dehydration is an atypical reaction for CYPs, which generally catalyse monooxygenation, but several CYPs from humans and plants catalyse aldoxime dehydration (44– 48). In the reaction, the reduction of heme iron Fe(III) to Fe(II) in the heme of CYPs enables the binding of the nitrogen atom of aldoximes to the heme iron, which allows charge transfer from Fe(II) to the aldoxime C=N bond, favouring elimination of the hydroxy group (44, 49, 50). CYP4GL4 and CYP30008A2 belong to different clans and share only 20% identity at the amino acid level (Fig. 3-A), indicating that the two nitrile-producing enzyme genes were recruited from non-paralogous genes in millipedes. The *K*_m_ value of CYP30008A2 towards (*E/Z*)-PAOx was 164 µM, 20 times higher than the 8.13 µM of CYP4GL4 (Fig. S13). However, the *K*_m_ value of CYP30008A2 is presumably low enough to catalyse the conversion of (*E/Z*)-PAOx into PAN in millipedes because the *K*_m_ of CYP30008A2 was not high compared to the 3.9–3200 µM of plant aldoxime-metabolising CYPs (48, 51–54).

The two CYPs exhibited acted on different substrates, not only L-Phe-derived (*E/Z*)-PAOx but also L-tryptophane-derived (*E/Z*)-IAOx and L-tyrosine-derived (*E/Z*)-4HPAOx (Fig. S14). CYP3201B1, which catalyses the conversion of PAN into (*R*)-MAN, also has broad substrate specificity (18). Thus, (*R*)-MAN biosynthesis in *C. hualienensis* is driven by the narrow substrate specificity of ChuaMOxS, which is responsible for the initial aldoxime production step (Fig. 4-E), whereas the enzymes responsible for subsequent reactions act on multiple amino acid-derived aldoximes and nitriles. The genes encoding both enzymes are similarly expressed in paraterga-containing defensive glands; therefore, the contribution of each enzyme to the nitrile-synthesising step is unclear. However, both enzymes could be involved in the nitrile-producing step, considering that the other two cyanogenic millipedes, *H. holstii* and *N. nodulosa*, harboured only one of the enzyme-encoding genes.

Cyanogenesis-related enzyme genes were specifically found in Polydesmida but not other millipede orders (Fig. 5). Furthermore, the enzyme clusters within arthropod enzymes exhibited no similarity or phylogenetic relationships with cyanogenesis-related enzymes from cyanogenic plants and insects (Figs. S10 and S16). These results indicate that Polydesmida millipedes independently evolved cyanogenesis-related enzyme genes from other cyanogenic organisms, ruling out other scenarios, such as divergent evolution or horizontal gene transfer from cyanogenic plants and the burnet moth. The independent evolution of identical enzyme activities can be either “convergent evolution” or “parallel evolution” (55). “Parallel evolution” is used when ancestral descendants possessing distinct biochemical activities but a shared structural lineage contemporarily evolve to synthesise the same metabolite. When distinct protein structures sharing no structural similarity result in the synthesis of the same metabolite, the term “convergent evolution” is employed. Thus, millipede cyanohydrin biosynthetic enzymes are the case of both convergent and parallel evolution with plant and insect enzymes. The recruitment of FMO (MOxS) in millipedes and CYPs in higher plants and insects constitutes an example of convergent evolution, whereas MOxS and fern FOS result from parallel evolution. Cyanohydrin synthesising steps are separated into two enzymes in millipedes. Generally, higher plants and insects synthesise cyanohydrin from aldoxime by a single CYP; sugar gum (*Eucalyptus cladocalyx*) utilises CYP706C55 and CYP71B103 (56), similar to the millipede (Fig. 1-B). The cyanohydrin synthesising step results from parallel evolution with millipedes and sugar gum. Thus, the cyanogenesis-related enzymes emerged multiple times independently through convergent and/or parallel evolution across arthropod and plant lineages. Our results will help to understand the mechanisms underlying the evolution of metabolic pathways among arthropods and plants.

## Materials and Methods

### Animals

*C. hualienensis* was collected from Kagoshima Prefecture, Japan, in November 2017 and 2018. The millipedes were reared in litter derived from the Japanese cedar *Cryptomeria japonica* D. Don at 22°C until use, as described previously (18).

### DNA preparation and genome sequencing

Genomic DNA was prepared from male millipedes for library construction. DNA samples were shipped to the Beijing Genomics Institute, Shenzhen, China, and sequenced using Illumina and PacBio Sequel sequencers. For detailed information about hybrid assembly, annotation, and other bioinformatic analysis, see SI Text.

### Phylogenetic analysis and divergence time estimation

A comparative genome sequence analysis of *C. hualienensis* was performed using 14 genomes. Orthologous groups were identified using Orthofinder2 (v. 2.5.4) with the ‘-M msa -T raxml-ng’ flag. Divergence time was estimated based on single-copy orthologues using BEAST. The calibration points used in BEAST were obtained from the TimeTree database (http://www.timetree.org/): centipedes versus millipedes (median time: 500 Mya), *Tribolium castaneum* and *Drosophila melanogaster* (median time: 334 Mya).

### Purification of ChuaHNL and its paralogous proteins with HNL activity

*P. pastoris* transformants were inoculated into 5 mL of YPD (1% yeast extract, 2% peptone, and 2% D-glucose). After 24 h culture at 30°C with shaking at 300 rpm, cells were harvested via centrifugation at 3,000 × *g* and 4°C for 10 min, resuspended in 500 mL of BMGH in a 2-L baffled flask, and grown at 30°C with shaking at 150 rpm for 14–18 h. Then, cells were harvested via centrifugation at 3,000 × *g* and 4°C for 10 min and resuspended in 500 mL of buffered minimal methanol medium, and 1% methanol was added as an inducer every 24 h. After 5 days of culture at 28°C with shaking at 150 rpm, the culture was centrifuged at 8,000 × *g* and 4°C for 15 min. The supernatant was recovered, and the pH was adjusted to 7.5 by adding aqueous 1 M K_2_HPO_4_. Resultant insoluble materials were removed via centrifugation at 13,000 × *g* and 4°C for 30 min and directly applied to a Ni Sepharose 6 fast flow column (Cytiva, Little Chalfont, UK), which was equilibrated with 20 mM KPB, pH 7.5, containing 300 mM NaCl and 20 mM imidazole. The column was washed with the same buffer, and the absorbed proteins were eluted using a linear imidazole gradient (20–500 mM) in the same buffer. The active fractions were concentrated and desalted using an Amicon Ultra-15 centrifugal filter device with a 10,000 nominal molecular weight limit cutoff (Merck Millipore, Billerica, MA, USA). It was loaded onto a Mono Q 5/50 column (Cytiva) equilibrated with 20 mM tricine-NaOH (pH 8.5) and eluted using a linear gradient of NaCl (0–500 mM) in the same buffer. The active fractions were pooled and concentrated using a centrifugal filtration device (Amicon Ultra 0.5 Centrifugal Filter Unit with ultracel-10 membrane; Merck Millipore).

### HNL assay

Enzyme activities for (*R*)-MAN synthesis and MAN cleavage were assayed using high-performance liquid chromatography and spectrophotometry, respectively. HNL activity during (*R*)-MAN synthesis was measured as previously reported (23), with slight modifications. In brief, the reaction (total volume = 0.25 mL) was initiated by adding potassium cyanide (100 mM) in citrate buffer (400 mM, pH 4.0) containing benzaldehyde (50 mM; 12.5 μL of 1 M benzaldehyde dissolved in dimethyl sulfoxide), and enzyme solution (0.1–1.0 U/mL). The final dimethyl sulfoxide concentration in the mixture was 5% (v/v). The mixture was incubated at 25°C for 5 min, and then aliquots (50 μL) of reactant were mixed with a nine-fold volume (450 µL) of a *n*-hexane:2-propanol (85:15) mixture containing 5 mM *p*-xylyl cyanide as an internal standard.

Finally, the organic phase was analysed using a high-performance liquid chromatography instrument (UFLC Prominence Liquid Chromatograph LC-20AD, Shimadzu, Kyoto, Japan) equipped with a chiral column (CHIRALCEL OJ-H, Daicel, Osaka, Japan; 250 mm length × 4.6 mm inner diameter (i.d.), particle size 5 µm). The amount of (*R*)-MAN was estimated using a standard curve obtained from the peak areas and concentration ratios of the authentic compounds and the internal standard. One unit of synthesis activity was defined as the amount of the enzyme that synthesises 1 μmol of (*R*)-MAN from benzaldehyde and potassium cyanide per minute.

For the MAN cleavage activity assay, the reaction was initiated by adding the enzyme sample to a reaction mixture (1 mL) consisting of (*R*,*S*)-MAN (10 mM) in citrate buffer (100 mM, pH 5.0). The reaction mixture contained 5% (v/v) dimethyl sulfoxide at the final concentration. The reaction velocity of benzaldehyde formation was monitored using an Evolution 201 ultraviolet-visible spectrophotometer (Thermo Fisher Scientific) at 25°C and 280 nm for 1 min (extinction coefficient of benzaldehyde = 1,352 L mol^−1^ cm^−1^) (23). One unit of cleavage activity was defined as the amount of enzyme that produces 1 μmol of benzaldehyde from (*R*,*S*)-MAN per minute.

### Collection of ChuaCYP sequences from myriapod genomes

CYPs from *C. hualienensis* and genome-sequenced myriapods were searched via BLASTP using CYP3201B1 (CYP2 clan, BAV93938.1), CYP3193A1 (CYP3 clan, BAV93917.1), CYP4GQ1 (CYP4 clan, BAV93949.1), CYP20A1, (CYP20 clan, BAV93936.1), and CYP302A1 (mitochondrial clan, BAV93914.1) from *C. hualienensis*. The *E*-value threshold was 1.0 × 10^−10^. CYPs with sequences that were too short (<400 aa) or too long (>600 aa) were eliminated from the dataset as putative malfunctioning proteins because animal CYPs are 500 amino acids in length (57). CYPs from *S. martima* were derived from a previous study (25).

### Heterologous production of ChuaCYPs in *S. cerevisiae* and identification of aldoxime- and nitrile-producing CYPs

Expression vectors were constructed to co-express cDNAs encoding ChuaCPR and ChuaCYPs. cDNAs encoding ChuaCPR (CHUA_001042) and ChuaCYPs were amplified via PCR and cloned into the pYeDP60 vector (58) using the NEBuilder HiFi DNA Assembly Master Mix (New England Biolabs, Ipswich, MA, USA). The oligonucleotide primers for PCR are summarised in Table S8. The inserted DNA sequences were confirmed via Sanger sequencing. ChuaCPR with the GAL10-CYC1 promoter and PGK terminator region was amplified via PCR, and the amplicon was cloned into the SphI site of pAUR101 (Takara) to generate pAUR-ChuaCPR. The plasmid was linearised using StuI and integrated into the genome DNA of *S. cerevisiae* INVScI (Thermo Fisher Scientific) through homologous recombination. The transformants were selected on a YPD-agar plate containing 0.4 µg/mL of aureobasidin A at 30°C for 3 days. Genome integration of ChuaCPR was confirmed through PCR, and the selected strain was named ChCR11 for subsequent experiments.

pYeDP60 vectors carrying cDNAs encoding CYPs were used to transform ChCR11. The transformants were selected on SGI-agar plates (0.67% yeast nitrogen base without amino acids, 0.1% casamino acid, 40 µg/mL L-tryptophan, 2% D-glucose, and 1.5% agar) at 30°C for 2 days. Transformants were inoculated into 0.5 mL of SGI medium (0.67% yeast nitrogen base without amino acids, 0.1% casamino acid, 40 µg/mL L-tryptophan, and 2% D-glucose) and cultured at 30°C for 24 h. Cells were harvested from 0.1 mL culture and resuspended in 0.5 mL of 2×SLI medium (1.34% (w/v) yeast nitrogen base without amino acids, 0.2% (w/v) casamino acid, 80 µg/mL L-tryptophan, and 4% (w/v) galactose) supplemented with 0.2% (w/v) raffinose. After culturing at 30°C for 24 h, the cells were harvested and resuspended in 0.2 mL of 2 mM L-Phe, 1 mM (*E/Z*)-PAOx, or 1 mM PAN in 50 mM KPB (pH 7.0) containing 20 mM glucose. The reaction was performed at 30°C with shaking at 1,200 rpm for 180 min.

The production of (*E/Z*)-PAOx from L-Phe was analysed using liquid chromatography–mass spectrometry (LC–MS) (Nexera HPLC coupled with an LCMS-2020, Shimadzu), equipped with a COSMOSIL 3C18-MS-II packed column (100 mm × 2.0 mm i.d., particle size 3 µm; Nacalai Tesque, Kyoto, Japan). The separation conditions were as follows: column oven temperature, 40°C; mobile phase A, 0.1% formic acid in water; mobile phase B, acetonitrile; 10–60% linear gradient of B for 7.5 min and 98% B for 2.5 min, delivered at 0.4 mL/min. MS was simultaneously performed in the negative-ion mode using an LCMS-2020 apparatus (Shimadzu) via electrospray ionisation. (*E/Z*)-PAOx ionised in the positive-ion mode was monitored in the extracted ion *m/z* 136 [M+H]^+^. The production of PAN from (*E/Z*)-PAOx and MAN from PAN was analysed using an ACQUITY UPLC H-Class system (Waters, Milford, MA, USA) equipped with a COSMOCORE 2.6 C_18_ column (50 mm × 2.1 mm i.d., particle size 2.6 µm; Nacalai Tesque) under the following conditions: column oven temperature, 40°C; mobile phase A, 0.1% formic acid in water; mobile phase B, acetonitrile; 10–60% linear gradient of B for 4 min and 60% B for 0.5 min, delivered at 0.4 mL/min. Aldoximes and nitriles were quantified at 210 nm.

### Preparation of microsomes harbouring CYP4GL4 and CYP30008A2 from yeast

Microsomes harbouring CYP4GL4 (CHUA_008638) and CYP30008A2 (CHUA_014123) were prepared from *S. cerevisiae* ChCR11, following a previously described method (48). The yeast cells were cultured in 2 mL SGI medium at 30°C for 24 h. The culture was transferred to 50 mL SGI medium at 30°C for 16 h. Yeast cells were harvested via centrifugation at 5,000 × *g* and 4°C for 10 min and resuspended in 500 mL of 2×SLI medium supplemented with 0.2% (w/v) raffinose in a 2-L baffled flask at 30°C for 24 h. Yeast cells were harvested via centrifugation (5,000 × *g*, 10 min, 4°C), washed with 50 mM HEPES-NaOH (pH 7.6) containing 20% glycerol, and weighed to obtain the wet cell weight. The cells were then resuspended in 1 mL of 50 mM HEPES-NaOH (pH 7.6) containing 20% glycerol, 1 mM DTT, and an appropriate amount of proteinase inhibitor cocktail (complete Mini EDTA-Free; Merck KGaA, Darmstadt, Germany) per gram of wet cells. The cells were disrupted using a Multi-beads Shocker system (Yasui Kikai, Osaka, Japan) and glass beads (diameter 0.5 mm), as described previously (18). The cell debris and beads were precipitated via centrifugation (10,000 × *g*, 10 min, 4°C), and the supernatant was ultracentrifuged (150,000 × *g*, 60 min, 4°C) to precipitate the fraction of intact microsomes. The microsomes were resuspended in 50 mM HEPES-NaOH (pH 7.6), containing 20% glycerol and 1 mM DTT and stored at −80°C until further use. The protein concentration of the microsomes was determined using the TaKaRa Bradford Protein Assay Kit (Takara) with bovine serum albumin as the standard.

### Aldoxime dehydratase assay

CYP4GL4 and CYP30008A2 activity was evaluated at optimal pH and temperature. A 100 µL reaction mixture containing the microsomal fraction harbouring CYP4GL4 (5 mg/mL), 50 mM KPB (pH 8.0), 1 mM NADPH, and 1 mM aldoxime was incubated at 35°C for 30 min. After the pre-incubation at 35°C for 5 min, the reaction was started by adding the microsomes. A 100 µL reaction mixture containing the microsomal fraction harbouring CYP30008A2 (5 mg/mL), 50 mM KPB (pH 7.0), 1 mM NADPH, and 1 mM aldoxime was incubated at 40°C for 30 min. After pre-incubation at 40°C for 5 min, the reaction was initiated by adding microsomes. The reaction was terminated by adding 0.1 mL of 0.2% formic acid in 40% acetonitrile. The resulting insoluble materials were precipitated via centrifugation (21,500 × *g*, 10 min, 4°C). The supernatant was analysed using an ACQUITY UPLC H-Class system (Waters) equipped with a COSMOCORE 2.6 C_18_ column (50 mm × 2.1 mm i.d., particle size 2.6 µm; Nacalai Tesque) under the following conditions: column oven temperature, 40°C; mobile phase A, 0.1% formic acid in water; mobile phase B, acetonitrile; 10–60% linear gradient of B for 4 min and 60% B for 0.5 min, delivered at 0.4 mL/min. Aldoximes and nitriles were quantified at 210 nm. The nitriles were quantified using standard curves generated from authentic compounds (PAN, Tokyo Chemical Industry, Tokyo, Japan; 4-hydroxyphenylacetonitrile, Tokyo Chemical Industry; indole-3-acetonitrile, Sigma-Aldrich, St Louis, MO, USA). To determine *K*_m_ and *k*_cat_, the reactions were conducted under standard assay conditions using 10–2000 µM (*E/Z*)-PAOx. Aldoximes were chemically synthesised as previously described (48). *K*_m_ and *k*_cat_ were determined by curve fitting the data using the drc package (version 3.0-1) (59) in R (4.1.2) (60) and the Michaelis–Menten equation.

### Collection of FMO sequences from myriapod genomes

FMOs from *C. hualienensis* and genome-sequenced myriapods were searched using BLASTP using FMO1 from *Homo sapiens* (group B clade I, UPI0003EAECD6), SNO from *Tyria jacobaeae* (group B, clade II, Q8MP06), FOS1 from *Phlebodium aureum* (group B, clade IV, QNT35807), and SCO7468 from *S. coelicolor* (Q8CJJ9). The *E*-value threshold was 1.0 × 10^−10^. The FMOs with short (<350 aa) and long (>550 aa) sequences were eliminated from the dataset because group B FMOs were 420–520 amino acids in length (33). The transmembrane region and subcellular localisation were predicted using Phobius (61) and DeepLoc 2.1 (62), respectively.

### Identification of ChuaFMOs catalysing (*E/Z*)-PAOx formation in *E. coli*

Each *E. coli* BL21(DE3) transformant carrying pGro7 and ChuaFMO expression plasmid was inoculated into LB containing 1% (w/v) glucose, kanamycin (50 µg/mL), and chloramphenicol (34 µg/mL) and cultured overnight at 37°C. Ten microlitres of culture was transferred to 2 mL of a TB-based autoinduction medium containing 2 mg/mL l-arabinose and cultured at 37°C for 2 h and further cultured at 16°C for 22 h. The culture was centrifuged at 5,000 × *g* and 4°C for 15 min, and the supernatant was extracted twice with 1 mL of ethyl acetate. The organic layer was collected and evaporated under a nitrogen atmosphere. The samples were dissolved in 100 µL of methanol, and a portion of the solution (1 µL) was analysed using an Agilent 7890A GC equipped with an HP-5ms column (30 m × 0.25 mm i.d.; 0.25 μm film thickness, Agilent Technologies). The column oven was programmed from 60°C for 2 min, then at 10°C min^−1^ to 290°C, and maintained for 5 min. Helium was used as the carrier gas at a flow rate of 1.0 ml min^−1^. MS was performed simultaneously using a GC system coupled with an Agilent 5975C inert XL EI/CI MSD equipped with a triple-axis detector (Agilent Technologies). All mass spectra were acquired in electron impact mode (ionisation energy: 70 eV). (*E/Z*)-PAOx was identified from the mass spectra, retention times, and comparisons with authentic compounds.

### Substrate specificity of ChuaMOxS (CHUA_003298)

*E. coli* BL21(DE3) carrying pGro7 and p28-ChuaMOxS was inoculated into LB containing 1% (w/v) glucose, kanamycin (50 µg/mL), and chloramphenicol (34 µg/mL) and was cultured overnight at 37°C. Twenty microlitres of the culture was transferred to 2 mL of a TB-based autoinduction medium containing 2 mg/mL L-arabinose and cultured at 37°C for 2 h and further cultured at 16°C for 22 h. A portion (500 µL) of culture was mixed with an equal volume of 40% acetonitrile containing 0.2% formic acid and centrifuged at 21,400 × *g* and 4°C for 15 min. The supernatant was collected, and the presence of PAOx, 4-hydroxyPAOx, or IAOx in the medium was detected using an LC–MS system equipped with a COSMOSIL 3C18-EB packed column (100 × 2.0 mm i.d., particle size 3 µm; Nacalai Tesuque). The separation conditions were as follows: column oven temperature, 40°C; mobile phase A, 0.1% formic acid in water; mobile phase B, acetonitrile; 20–98% linear gradient of B for 7.5 min and 98% B for 2.5 min, delivered at 0.4 mL/min. MS was simultaneously performed in positive-ion mode, using an LCMS-2020 apparatus (Shimadzu), via electrospray ionisation (interface voltage, 3 kV; interface temperature, 300°C; DL temperature, 250°C; heat block temperature, 400°C; nebulising gas, 3 L/min; drying gas, 10 L/min; heating gas, 10 L/min). Aldoximes ionised in positive-ion mode were monitored in the extracted ions: PAOx, *m/z* 136 [M+H]^+^; 4HPAOx, *m/z* 152 [M+H]^+^; and IAOx, *m/z* 175 [M+H]^+^. The (*E/Z*)-PAOx accumulated in the medium was quantified using standard curves generated from authentic compounds.

### Phylogenetic analysis

Phylogenetic analyses of HNL, CYPs, and FMOs were performed. Multiple sequence alignments were performed using MAFFT software (63). A phylogenetic tree was constructed using the maximum-likelihood method and RAxML-NG (64) with the best-fit amino acid substitution model (HNL; WAG+I+G4+F, CYP; LG+I+G4+F, FMO; LG+G4+F) determined based on ModelTest-NG (65) and the Akaike Information Criterion. The tree was evaluated using bootstrap analysis with 500 (CYP) or 1000 (HNL and FMO) replicates.

### Statistical analysis

Statistical analysis was performed using the multcomp package (66) in R (version 4.12), and differences were analysed using Welch’s t-test or Tukey’s honest significant difference test. Differences were considered statistically significant at P < 0.05.

## Supporting information

SupportingInformation

## Data availability

The final assemblies were submitted to DDBJ under accession number BAAGAE01000000– BAAGAE010000055. The raw reads generated in this study were deposited to the DDBJ database under the BioProject accession number PRJDB13209.

## Acknowledgements

We thank Dr D. Nelson for naming the cytochrome CYPs from *C. hualienensis* and Ms. M. Fukutani for construction of *Pichia* expression plasmids. This work was supported by the JST ERATO Asano Active Enzyme Molecule Project (Grant Number JPMJER1102), Japan, a grant-in-aid for Scientific Research (S) and (A) from The Japan Society for Promotion of Sciences (Grant Numbers 17H06169 and 22H00361, respectively) to YA, and a Toyama Prefectural University Grant for Incentive Research to TY.

