## SupportingInformation for "Cyanogenic millipede genome illuminates convergent evolution of cyanogenesis-related enzymes"

### Supporting Information for
### Cyanogenic millipede genome illuminates convergent evolution of cyanogenesis-related enzymes

Takuya Yamaguchi[a, *], Yasuhisa Asano[a, *]

[a] Biotechnology Research Center and Department of Biotechnology, Toyama Prefectural University, 5180 Kurokawa, Imizu, Toyama 939-0398, Japan

* Yasuhisa Asano

* Takuya Yamaguchi

**This PDF file includes:**

Supporting text
Figures S1 to S21
Tables S1 to S10
SI References

#### Supporting text

##### Chemicals

(*E/Z*)-PAOx and (*E/Z*)-4-hydroxyphenylacetaldoxime (4HPAOx) were synthesized by condensation of hydroxylamine with phenylacetaldehyde and 4-hydroxyphenylacetaldehyde, respectively (1). 4-Hydroxyphenylacetaldehyde was prepared from L-tyrosine via sodium hypochlorite oxidation (2). (*E/Z*)-Indole-3-acetaldoxime (IAOx) was synthesized by condensation of hydroxylamine with the corresponding aldehyde derived from 2-(1*H*-indol-3-yl)ethanol (3). Other chemicals were purchased from commercial suppliers.

##### DNA preparation and genome sequencing

Genomic DNA was prepared from male millipedes for library construction. To avoid microbial contamination from the digestive system, the animals were degutted in phosphate-buffered saline before being immediately frozen in liquid nitrogen for storage at −80°C until use. Degutted millipedes were ground in liquid nitrogen using a mortar and pestle. Genomic DNA was purified using the Blood & Cell Culture DNA Maxi Kit (Qiagen, Valencia, CA, USA). DNA samples were shipped to the Beijing Genomics Institute, Shenzhen, China, and sequenced using Illumina and PacBio Sequel sequencers.

##### Genome size estimation by *k*-mer analysis

To estimate the genome size of *C. hualienensis*, *k*-mers in sequencing reads (Illumina short reads, 270 bp library) were counted using jellyfish 2.2.10 (4). A frequency histogram of the 21-mers was obtained and analysed using GenomeScope 1.0 (5).

##### Genome assembly

First, mitochondrial genome-derived reads were removed from the raw read sequences. Illumina and PacBio reads were mapped to the assembled mitochondrial genome using BWA-MEM 0.7.17 (6) and Mimimap2.1, respectively (7). The mitochondrial genome-filtered Illumina paired-end reads were assembled using Platanus 1.2.4 (8). The resultant contigs and PacBio Sequel reads longer than 1 kb were applied to the hybrid assembly using the DBG2OLC and Sparc software pipelines (9). The PacBio Sequel reads were mapped to contigs using minimap and polished twice using Racon. Next, 300 × paired-end reads were mapped to contigs using BWA-MEM, and the contigs were polished twice using Pilon (10). Finally, 33 × mate-paired reads were mapped to polished contigs, which were scaffolded using BESST (11). Scaffold completeness was assessed using the BUSCO (12) and Arthropoda dataset odb9 (13) .

##### RNA preparation and RNA sequencing

Total RNA was prepared from antennae from male and female, segments with or without defensive glands, and gut using the RNeasy mini Kit (Qiagen). Purified total RNA was sequenced at the Beijing Genomics Institute using a Novaseq 6000 (Illumina, Hayward, CA, USA).

##### RNA-seq assembly

RNA-seq data (Table S9 and S10) of *C. hualienensis* were assembled using the Trinity-V2.8.4 (14) with --jaccard_clip option.

**Annotation of repetitive sequences**

De novo and homology-based approaches were integrated to search for transposable elements and annotate repetitive sequences. RepeatModeler version 2.0.1 (15) was run on the unannotated assembly to identify and classify de novo repeat families. RepeatModeler employs two de novo repeat-finding programs, RECON version 1.08 and RepeatScout version 1.0.5, to identify repeat element boundaries and build consensus models of putative interspersed repeats. The sequence was aligned to GenBank's non-redundant protein database using diamond BLASTX with an E-value cutoff of $1 \times 10^{-5}$ (--sensitive) to ensure that repeat sequences in the library did not contain large families of protein-coding genes that are not transposable elements. Repeat masking was performed on the assembled genome using RepeatMasker version 4.0.9 (http://www.repeatmasker.org/) against repetitive sequences in the RepeatMasker consensus library (20150807; www.girinst.org) and a custom species-specific repeat library generated using RepeatModeler.

**Gene prediction and annotation**

To predict the genes, we employed the funannotate pipeline (https://github.com/nextgenusfs/funannotate). The pipeline utilises several ab initio gene predictors, including Augustus, GeneMark-ET v4.46, snap, glimmerHMM, and the PASA annotation pipeline. Augustus and GeneMark-ET were pretrained using BRAKER software. RNA-seq reads (Table S9) were mapped to scaffolds using STAR v2.7.7a. Resultant BAM files were merged and applied to BRAKER software. De novo RNA-seq assembly of the millipede was applied to the funannotate package.

**Extraction and quantification of (*R*)-MAN from *C. hualienensis***

The millipedes were separated into male and female. Their body weight was weighed, and MAN was extracted with 200 µL of MeOH containing 1 mM PAN per 100 mg body weight. A portion (2 µL) of the extract was analyzed using an Nexera UPLC (Shimadzu, Kyoto, Japan) system equipped with a COSMOCORE 2.6$C_{18}$ column (50 mm × 2.1 mm i.d., particle size 2.6 µm; Nacalai Tesque, Kyoto, Japan) under the following conditions: column oven temperature of 40 °C; mobile phase A, 0.1% formic acid in water, mobile phase B, acetonitrile, and 10–60% linear gradient of B for 4 min and 60% B for 0.5 min delivered at 0.4 mL/min. The amount of (*R*)-MAN was estimated using the standard curve obtained from the peak area ratio and concentration ratio of the authentic compounds and the internal standard.

**Heterologous production of ChuaHNL and its paralogous proteins in *P. pastoris***

ChuaHNL and its paralogous proteins were heterologously produced as N-terminal His-tagged proteins as previously described (16). Signal peptides were predicted using the SignalP 6.0 server (https://services.healthtech.dtu.dk/services/SignalP-6.0/). Next, ChuaHNL paralogous protein-coding sequences without signal peptides were synthesised, and the codons were optimised for expression in *P.*

*pastoris* using GeneArt Strings (Thermo Fisher Scientific, Waltham, MA, USA). pPICZαA was linearised via inverse PCR using pPICZα-His-Syn-ChuaHNL as a template DNA (Zhai et al., 2019). The synthesised DNA fragments were cloned into a linearised vector using an In-Fusion HD Cloning Kit (Clontech Laboratories, Palo Alto, CA, USA) or the NEBuilder HiFi DNA Assembly Master Mix. The inserted DNA sequence was verified using Sanger sequencing.

The constructed vectors were linearised via digestion with SacI and transformed into *P. pastoris* PpPDI/GS115 cells (16) harbouring genomic DNA-integrated *AOX1::PpPDI*, using a Pichia EasyComp Transformation Kit (Thermo Fisher Scientific), according to the manufacturer's instructions. Transformants were selected on YPDS (1% yeast extract, 2% peptone, 2% D-glucose, and 1 M sorbitol) agar medium containing 100 µg/mL zeocine. His-tagged recombinant protein production was evaluated as follows. Selected *P. pastoris* transformants were inoculated into 2 mL of YPD and cultured at 30°C for 16 h. Then, 100 mL of buffered minimal glycerol medium [BMGH; 100 mM potassium phosphate buffer (KPB; pH 7.0), 1.34% yeast nitrogen base without amino acid, $4\times10^{-5}$% biotin, 0.004% L-histidine, and 1.0% glycerol] in a 500 mL baffled flask. The cells were harvested via centrifugation and resuspended in buffered minimal methanol medium (the same as BMGH, but 1% (v/v) methanol was added instead of 1% (v/v) glycerol) at $OD_{600}$ = 5). After 6 days culture at 28°C with shaking at 150 rpm, the culture was centrifuged at 8,000 × *g* and 4°C for 15 min. The supernatant was recovered, and the pH was adjusted to 7.2–7.5 by adding 1 M $K_2HPO_4$. Resultant insoluble materials were removed via centrifugation at 20,000 × *g* and 4°C for 15 min. To concentrate the His-tagged proteins secreted into the medium, the supernatant was applied to a column packed with 1 mL complete His-Tag Purification Resin (Roche Applied Science, Basel, Switzerland), which was equilibrated with 20 mM KPB (pH 7.5) containing 300 mM NaCl and 20 mM imidazole. The column was washed with the same buffer, and the absorbed proteins were eluted with 5 mL of the same buffer containing 300 mM imidazole.

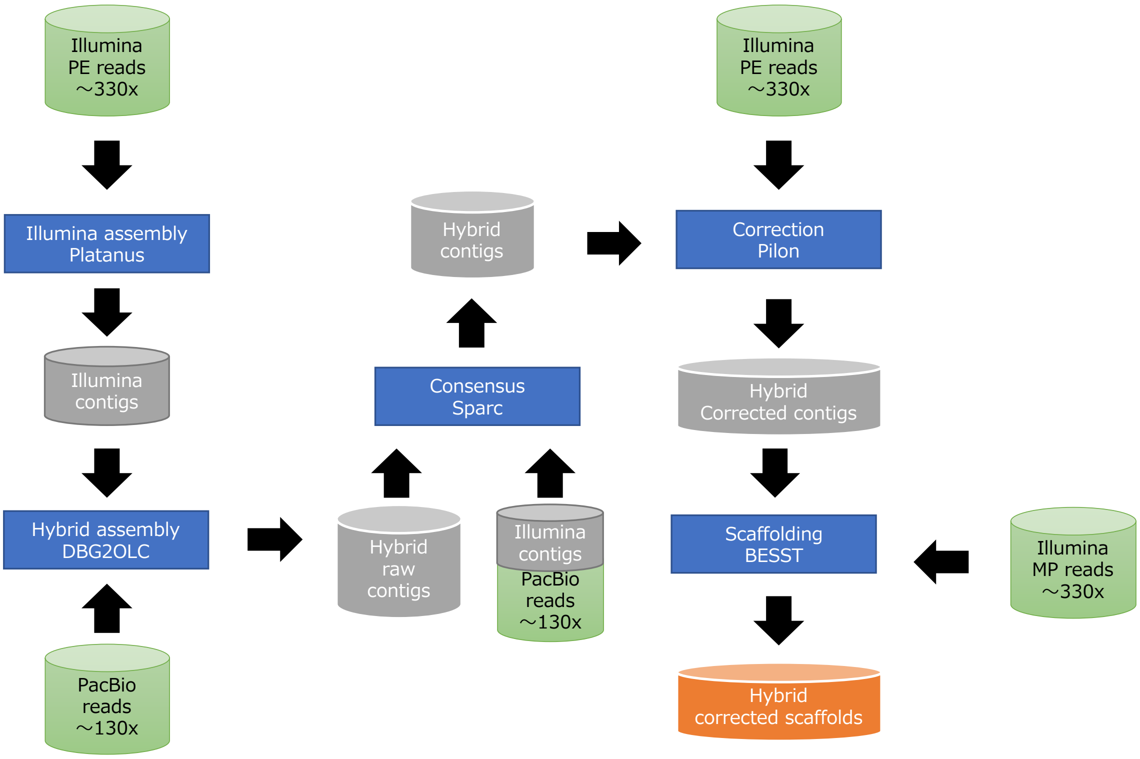

**Fig. S1. Overview of the processing pipeline used for the assembly of the *Chamberlinius hualienensis* genome (see materials and methods for detail).**

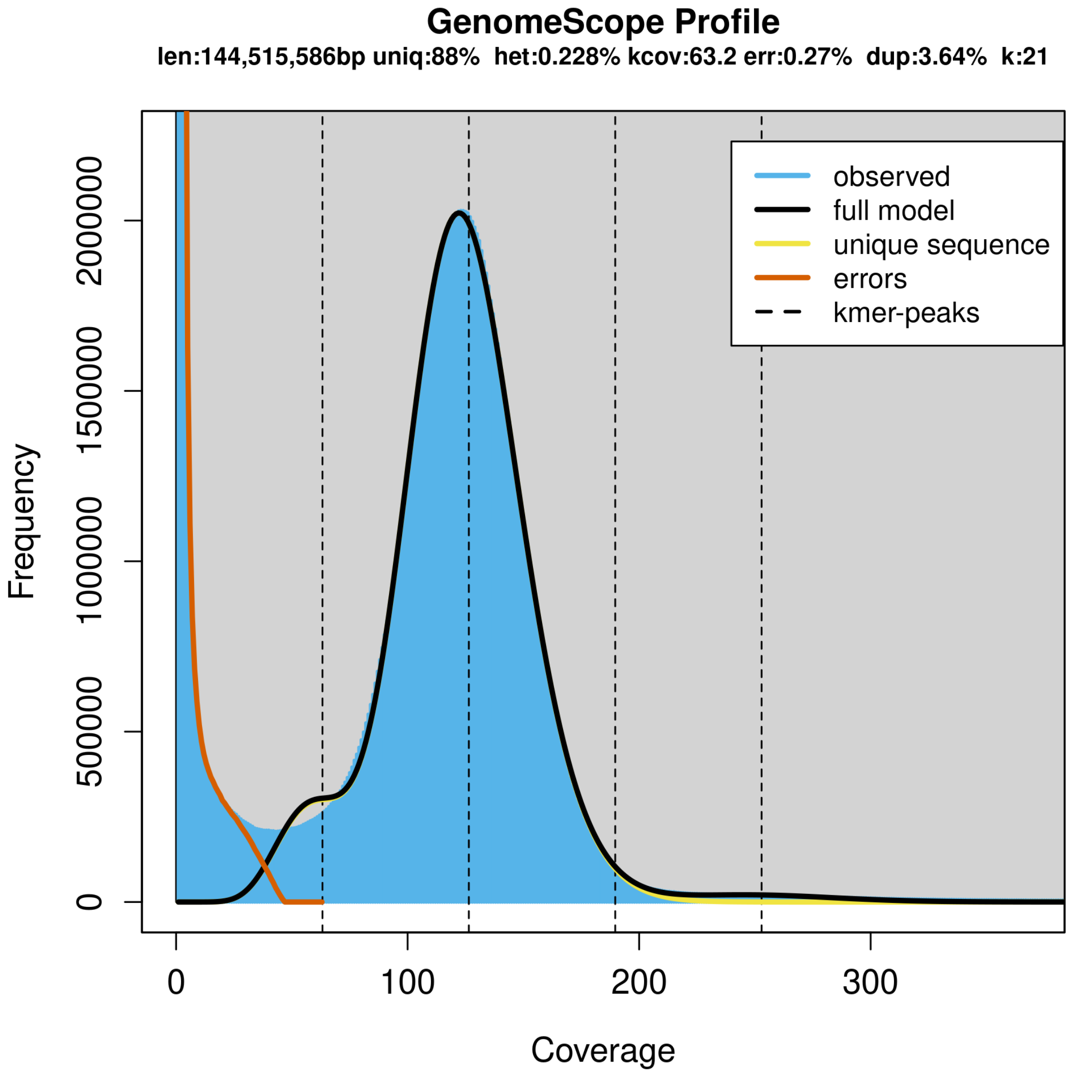

**Fig. S2. GenomeScope plot of the 21-mer content within the C. hualienensis genome.** Dataset show the fit of the GenomeScope model (black) based on 21-kmers in Illumina HiSeq sequence reads

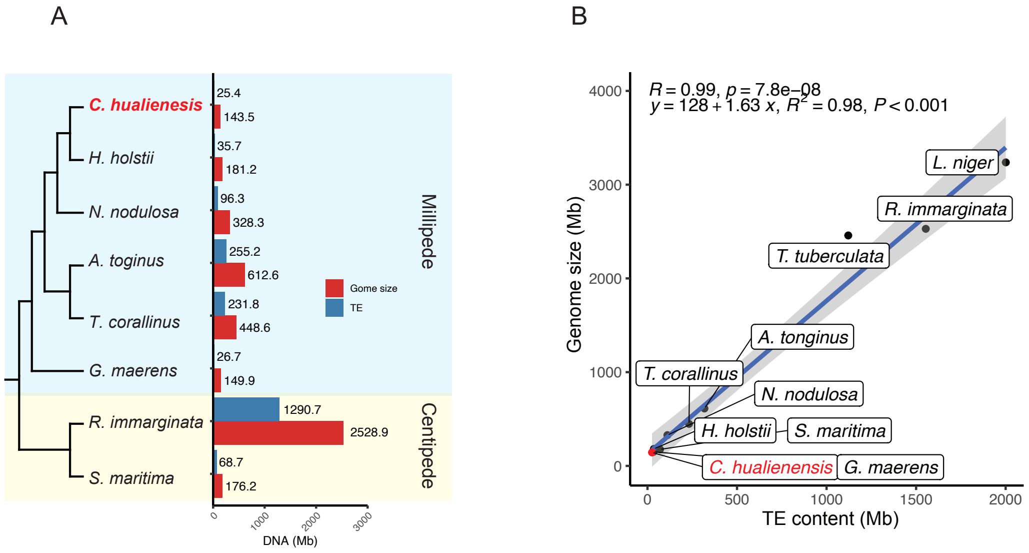

**Fig. S3. Correlations of genome sizes and TE contents among millipedes and centipedes**. A. Genome sizes and TE contents of millipedes and centipedes. B. Correlations of genome sizes and TE contents among millipedes and centipedes.

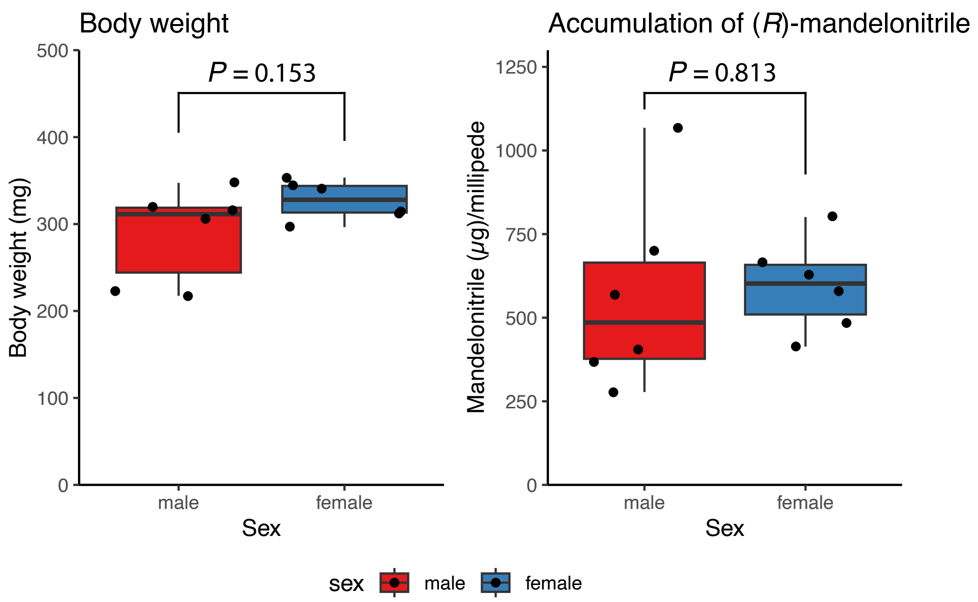

**Fig. S4. Body weight and accumulation of (*R*)-mandelonitrile in *Chamberlinius hualienensis*.** Each boxplot shows median (the center horizontal), interquartile range (upper and lower edges of the box), and 1.5 times the interquartile range (whisker) (n = 6). Welch's *t* test was used to analyze the data for significant differences between male and female. *P* values less than 0.05 were considered statistically significant.

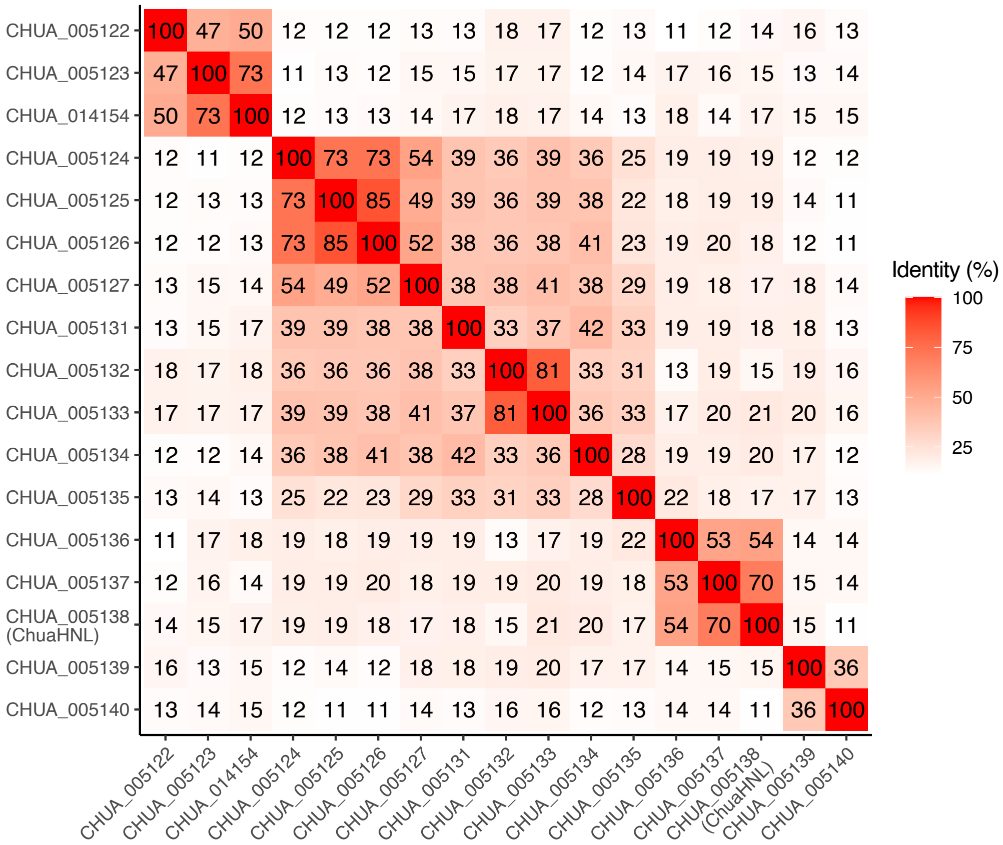

**Fig. S5. Amino acid sequence identity matrix of ChuaHNL and its paralogous proteins.**

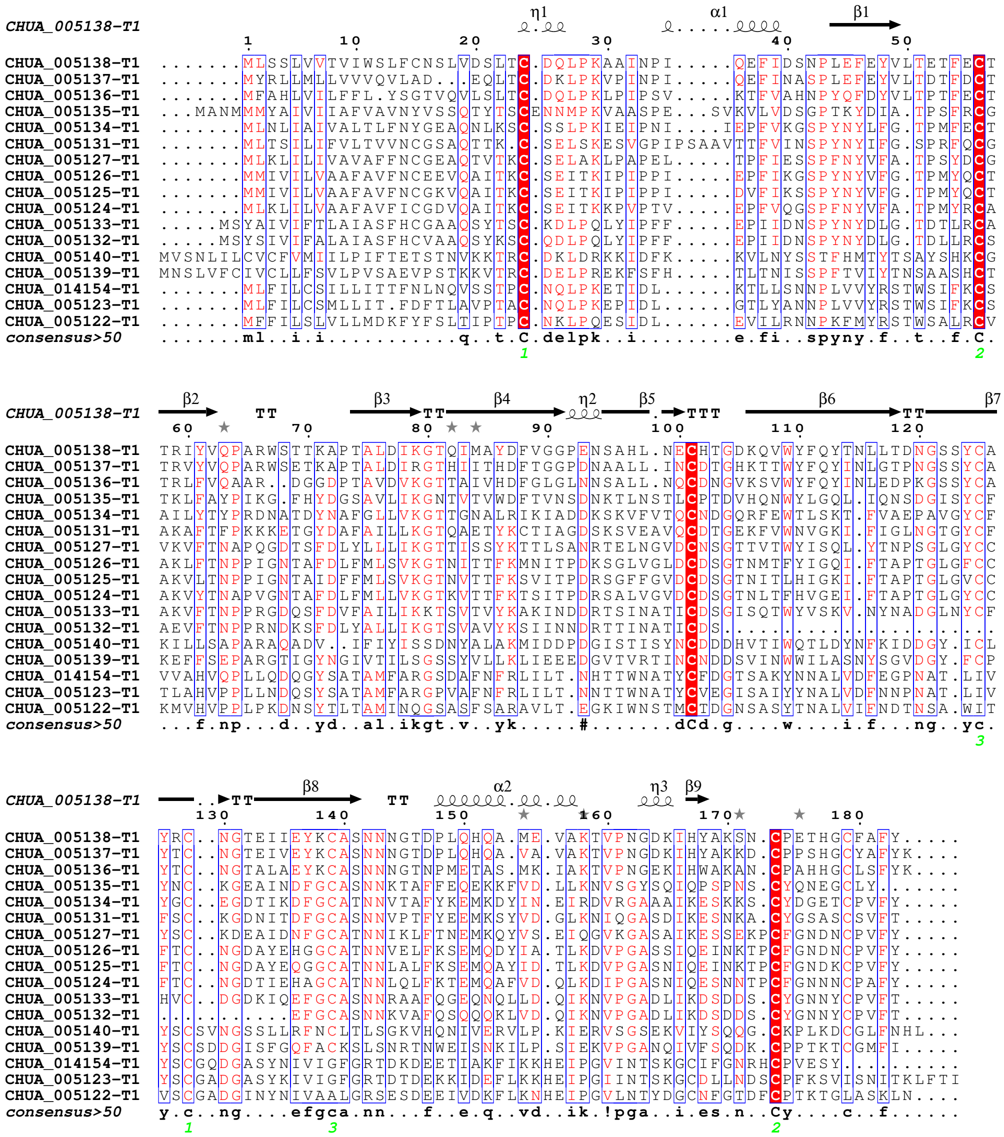

**Fig. S6. Amino acid sequence alignment of ChuaHNL and its paralogous proteins.**

A red background identifies strictly conserved residues; red letters indicate residues well conserved within a group, according to a Raisler matrix; and all other amino acids are shown in black. Residues conserved between groups are boxed with a blue line.

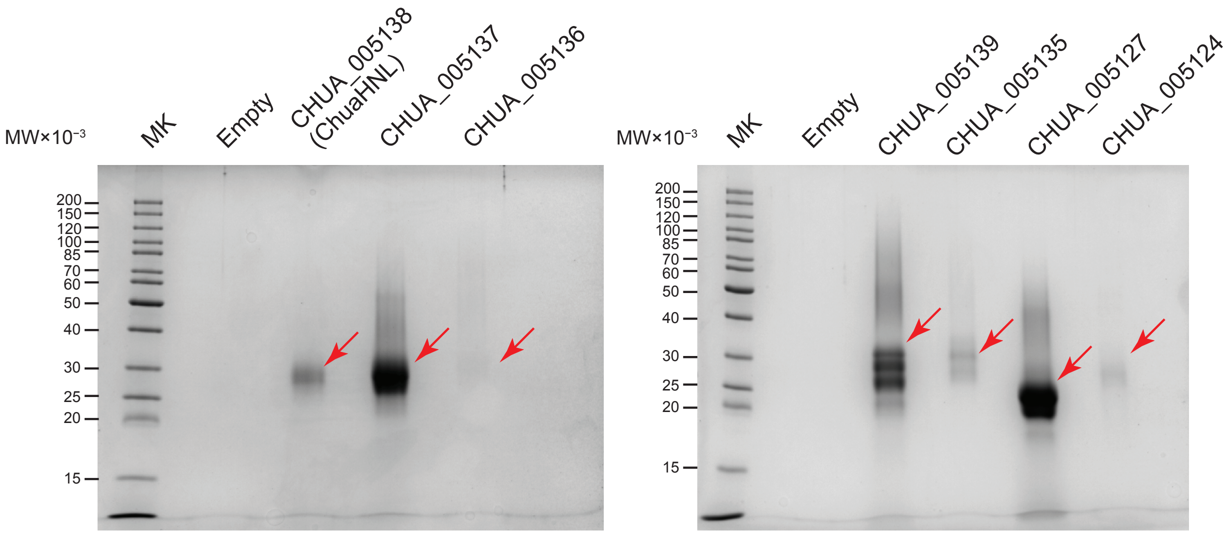

**Fig. S7. Heterologous production of ChuaHNL and its paralogous proteins in *Pichia pastoris*.**

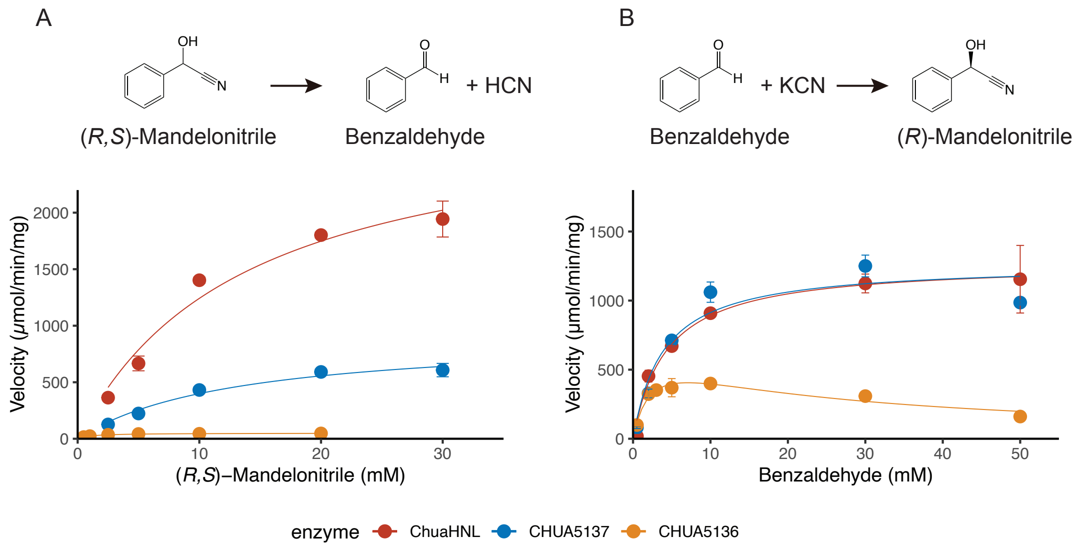

**Fig. S8. Substrate saturation curves for CHUA_005138 (ChuaHNL), CHUA_005137, and CHUA_005136 with (*R,S*)-mandelonitrile (mandelonitrile cleavage reaction) (A) or benzaldehyde ((*R*)-mandelonitrile synthetic reaction) (B).**

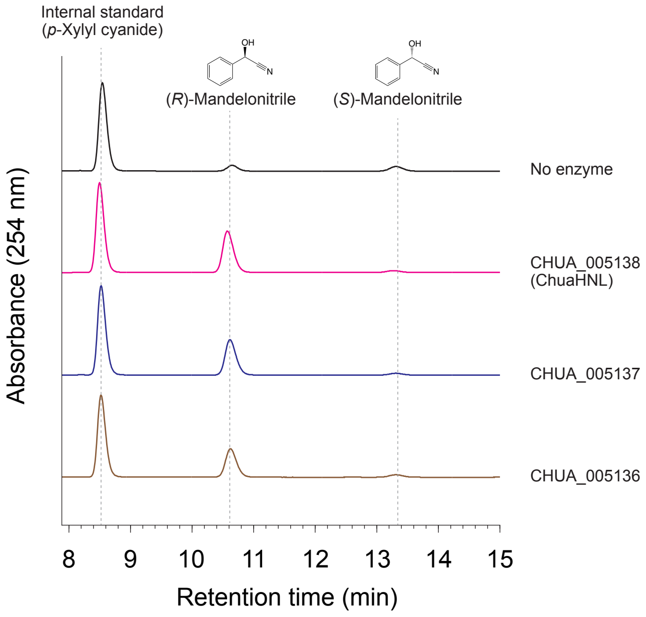

**Fig. S9. (*R*)-mandelonitrile synthetic reaction catalyzed by CHUA_005138 (ChuaHNL), CHUA_005137, and CHUA_005136.** The reaction products were extracted and analyzed by a high performance liquid chromatograph equipped with a chiral column.

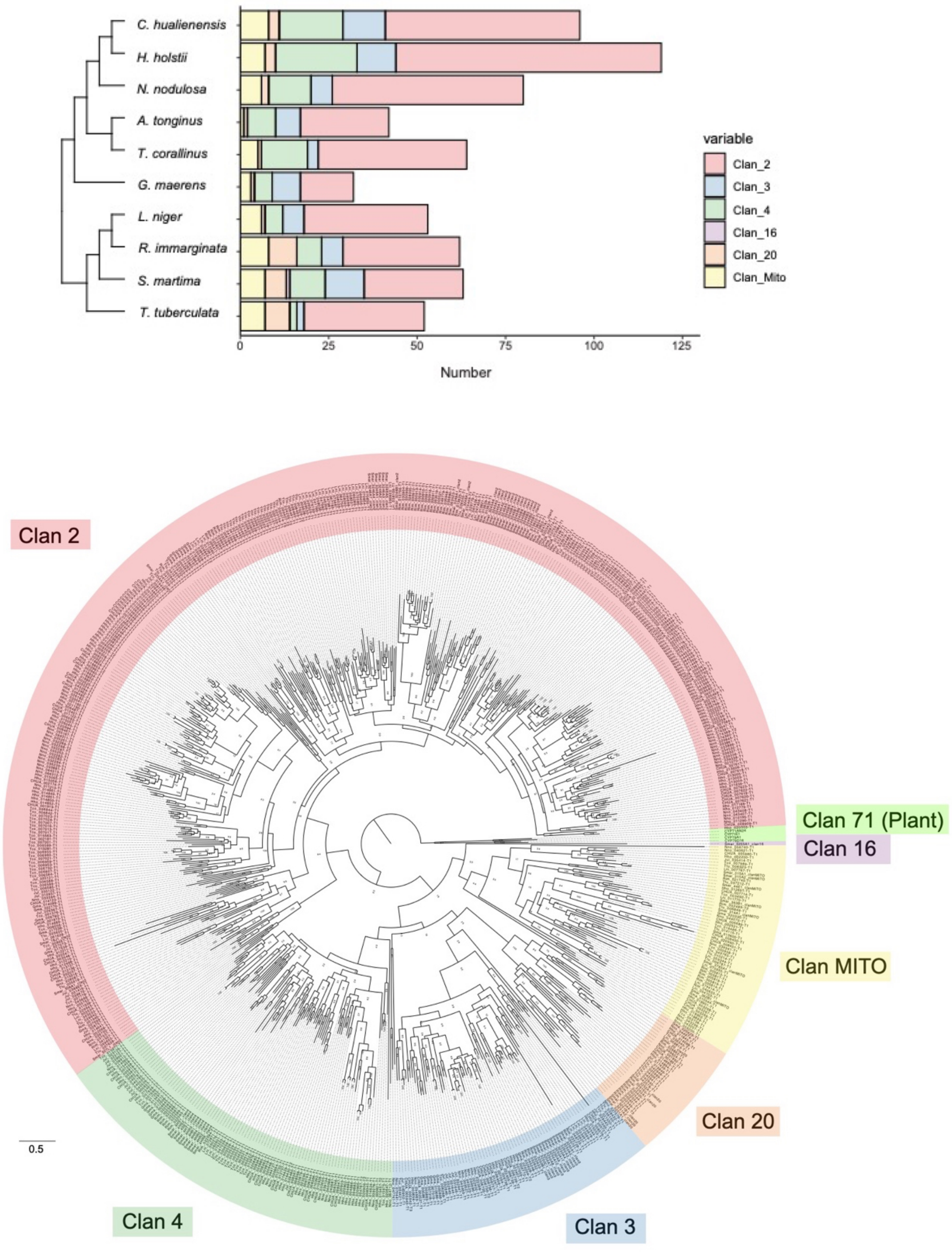

**Fig. S10. Phylogenetic tree of cytochrome P450s**

Number and phylogenetic tree of cytochrome P450s from Myriapoda species (*C. hualienensis*, *H. holstii*, *N. nodulosa*, *A. tonginus*, *T. corallinus*, *G. maerens*, *L. niger*, *R. immarginata*, *S. martima*, *T. tuberculata*). The bar indicates 50% divergence

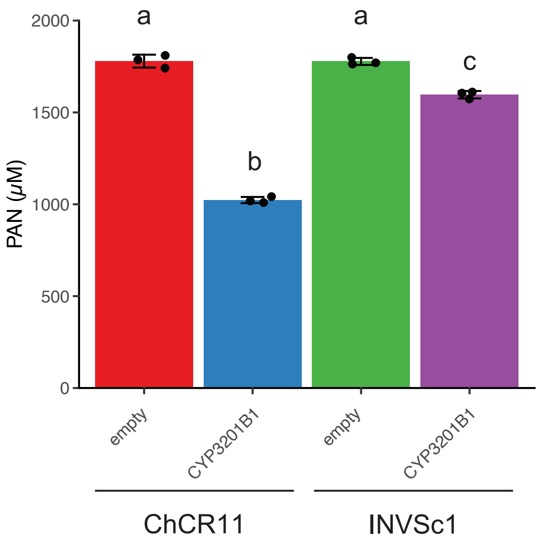

**Fig. S11. Comparison of phenylacetonitrile (PAN) metabolizing activity of *Saccharomyces cerevisiae* ChCR11 and INVSc1 expressing *CYP3201B1*.**

Yeast cells carrying the CYP3201B1 expression plasmid or an empty vector were incubated with PAN. PAN after the reaction was quantified using ultra-performance liquid chromatography. Reactions were carried out in triplicate ($n = 3$), error bars show the standard deviation of the replicate measurements, the error bar centers are the means of the replicate measurements, and the replicate measurements are represented as black dots. Bars labelled with different letters indicate a significant difference ($P < 0.05$) as determined by Tukey’s honest significant difference test.

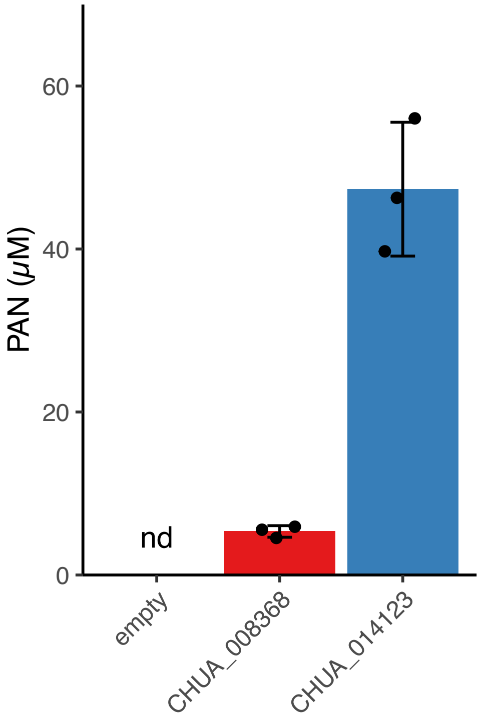

**Fig. S12. Formation of phenylacetonitrile (PAN) from (*E/Z*)-phenylacetaldoxime by yeast harboring CHUA_008368 (CYP4GL4) and CHUA_014123 (CYP30008A2).** Yeast cells carrying the CHUA_008368 and CHUA_014123 expression plasmid or an empty vector were incubated with (*E/Z*)-phenylacetaldoxime. The formation of PAN was quantified using ultra-performance liquid chromatography. Reactions were carried out in triplicate ($n = 3$), error bars show the standard deviation of the replicate measurements, the error bar centers are the means of the replicate measurements, and the replicate measurements are represented as black dots. nd, not detected.

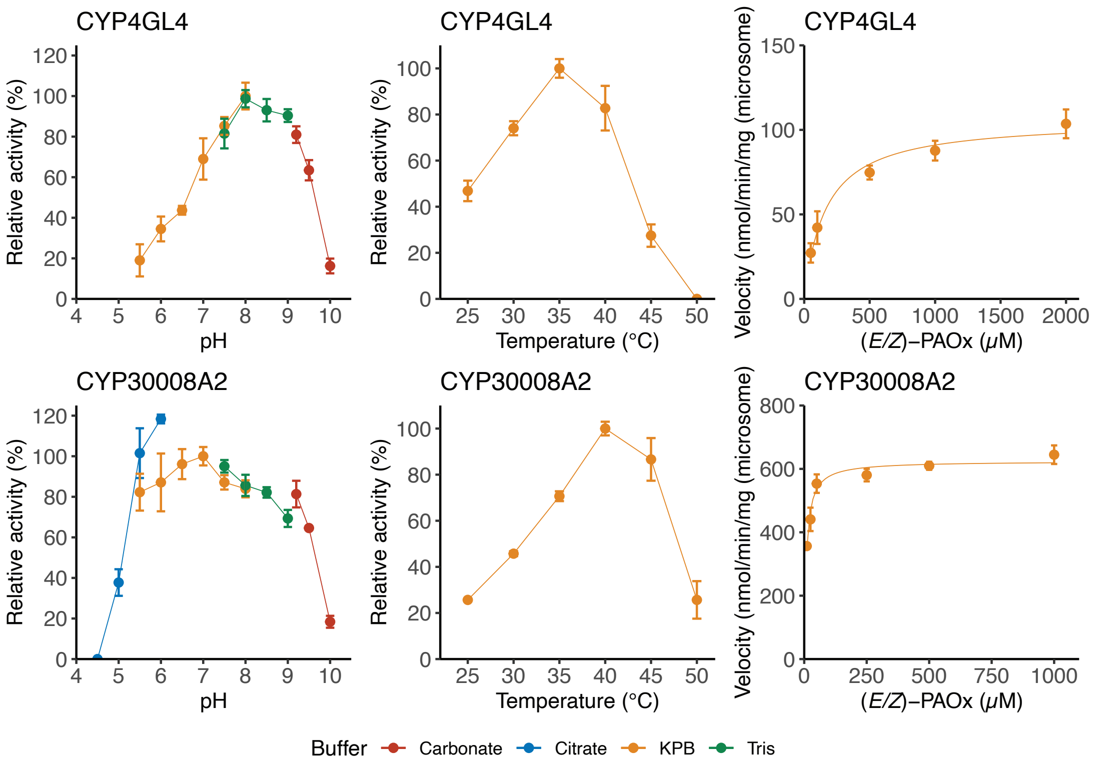

**Fig. S13. Characterization of CYP4GL4 (CHUA_008368) and CYP30008A2 (CHUA_014123) catalyzing dehydration of (*E/Z*)-phenylacetaldoxime (PAOx) into phenylacetonitrile.** Reactions were carried out in triplicate ($n = 3$), error bars show the standard deviation of the replicate measurements, and the error bar centers are the means of the replicate measurements.

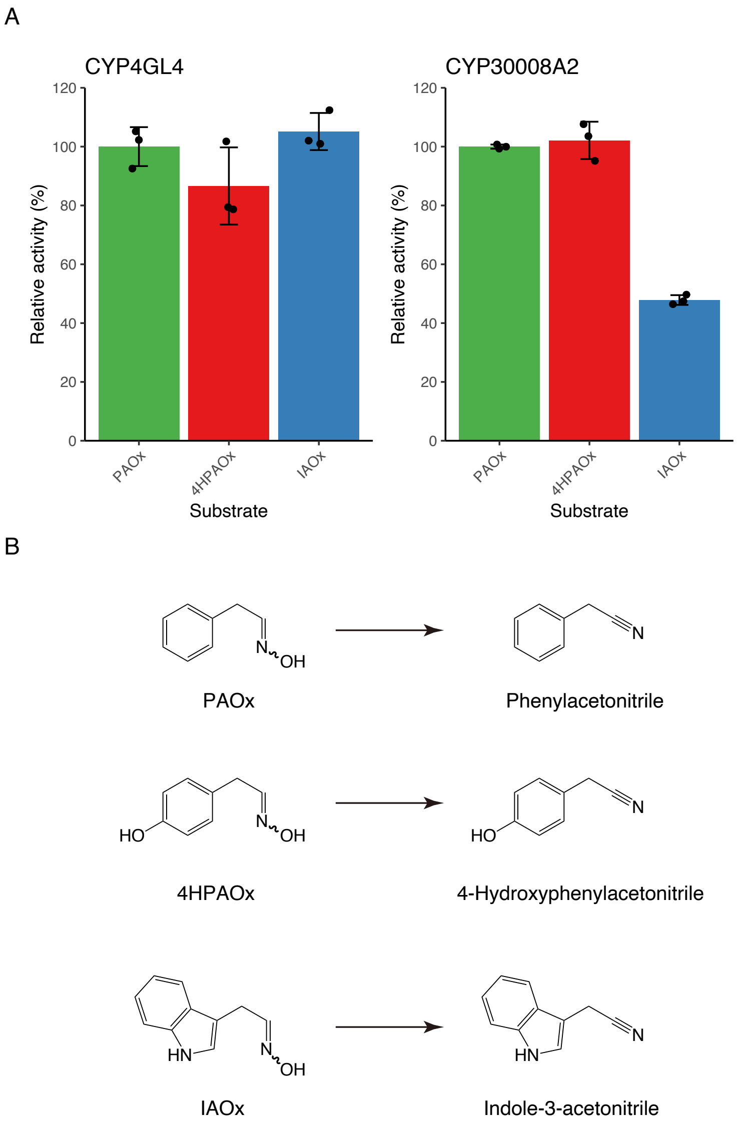

**Fig. S14. Substrate specificity of CYP4GL4 (CHUA_008368) and CYP30008A2 (CHUA_014123) toward aromatic aldoximes.** A. Microsome harboring CYP4GL4 and CYP3008A2 were incubated with 1 mM aldoximes in the presence of NADPH. Corresponding nitriles produced from (*E*/*Z*)-phenylacetaldoxime (PAOx), (*E*/*Z*)-4-hydroxyphenylacetaldoxime (4HPAOx), and (*E*/*Z*)-indole-3-acetaldoxime (IAOx) were quantified using ultra performance liquid chromatography. The activity toward PAOx was defined as 100%. Reactions were carried out in triplicate ($n = 3$), error bars show the standard deviation of the replicate measurements, the error bar centers are the means of the replicate measurements, and the replicate measurements are represented as black dots. B. The reactions catalyzed by the two enzymes.

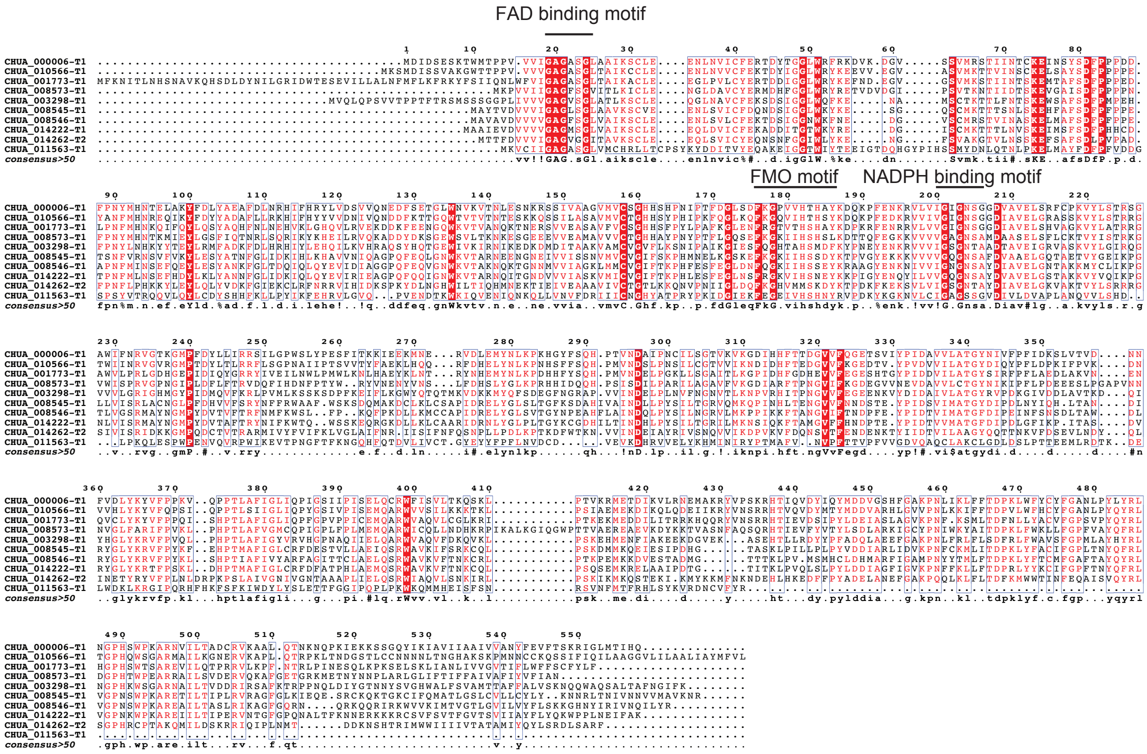

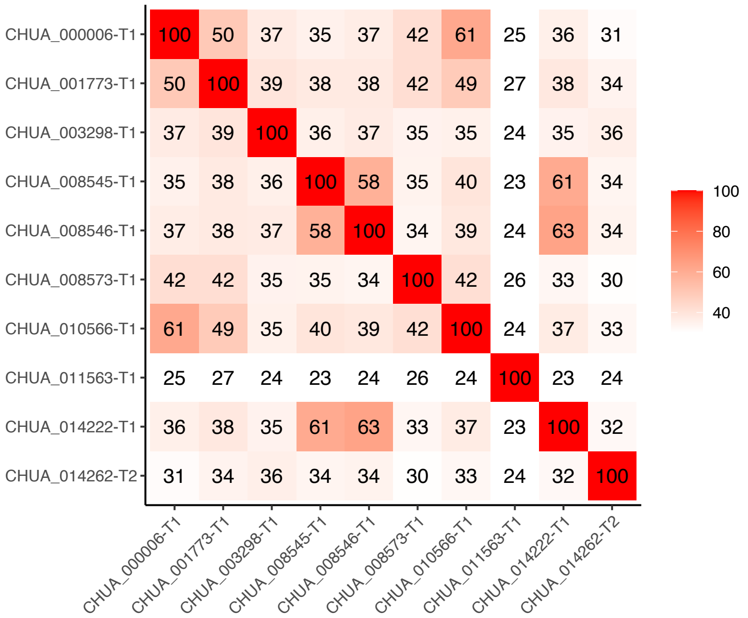

**Fig. S15. Amino acid sequence alignment and sequence identity matrix of *Chamberliniensis hualienensis* flavin-dependent monooxygenases.**

A red background identifies strictly conserved residues; red letters indicate residues well conserved within a group, according to a Raisler matrix; and all other amino acids are shown in black. Residues conserved between groups are boxed with a blue line.

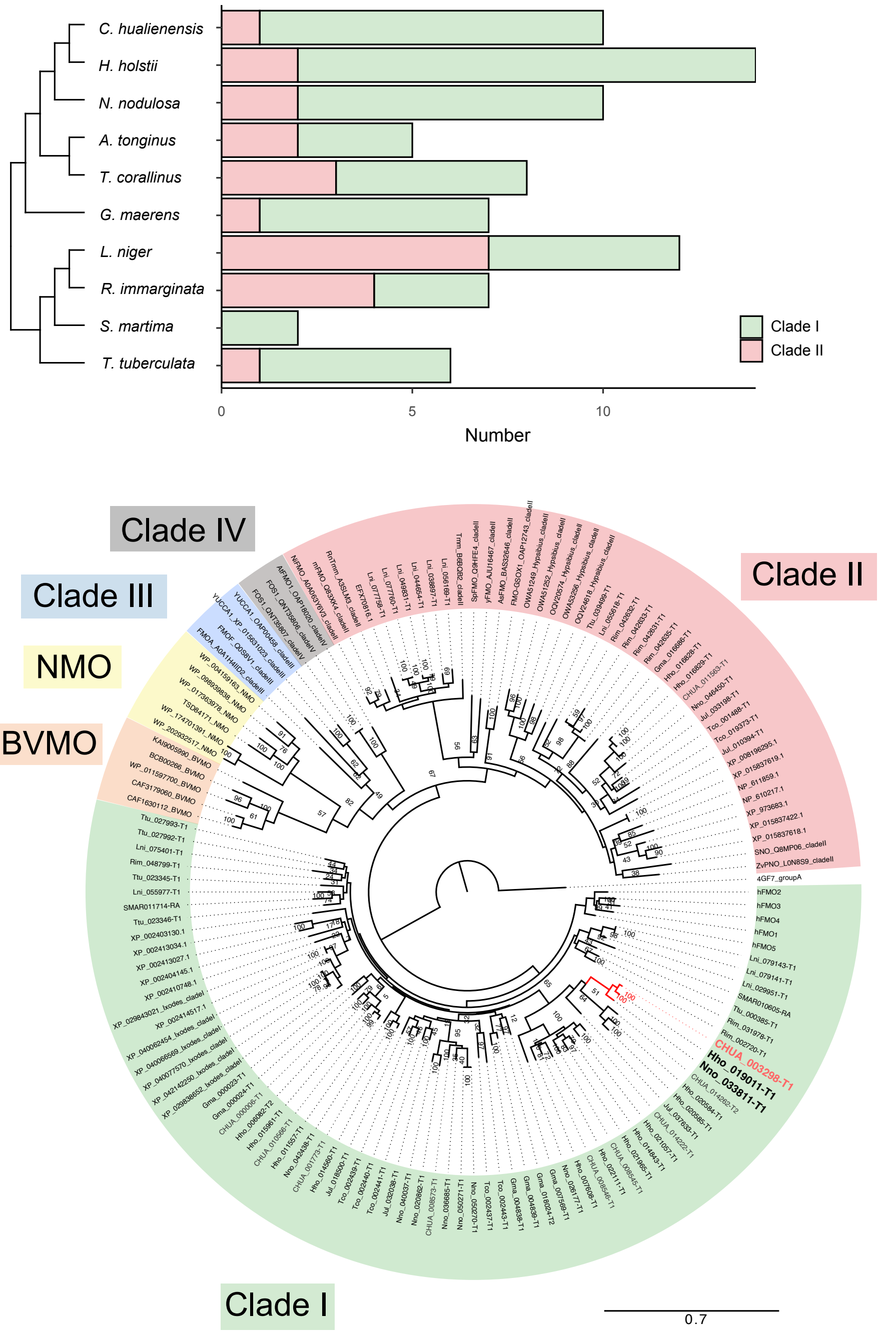

**Fig. S16. Flavin-dependent monooxygenases in Myriapoda**

Phylogenetic tree of flavin-dependent monooxygenases from Myriapoda species (*C. hualienensis*, *H. holstii*, *N. nodulosa*, *A. tonginus*, *T. corallinus*, *G. maerens*, *L. niger*, *R. immarginata*, *S. martima*, *T. tuberculata*). The bar indicates 70% divergence.

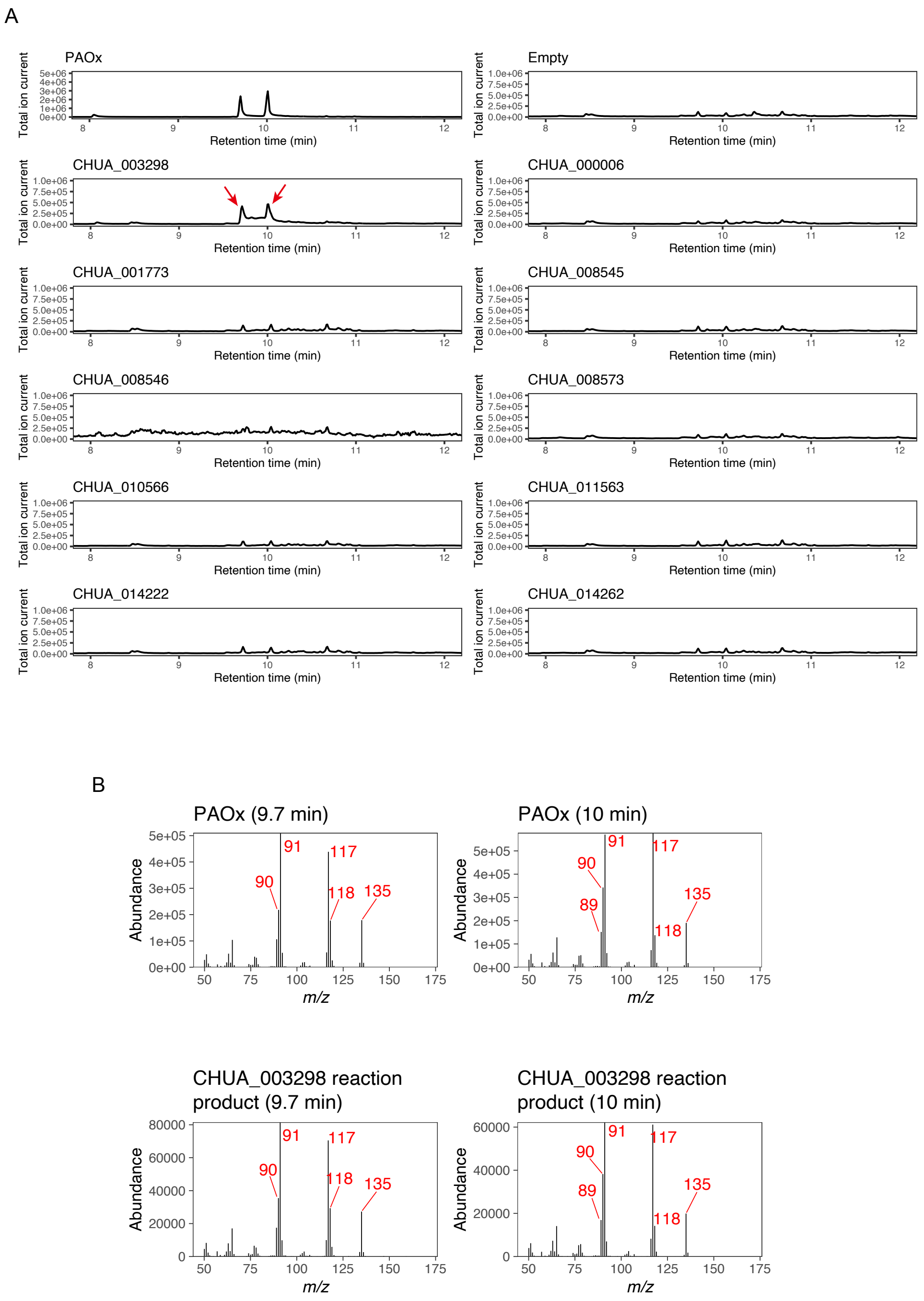

**Fig. S17. Identification of PAOx-producing flavin dependent monooxygenase from *Chamberlinius hualienensis*.** A. Identification of the (*E/Z*)-PAOx-producing FMOs. *Escherichia coli* cells carrying FMOs from *C. hualienensis* expression plasmids or an empty vector were cultured. The accumulation of (*E/Z*)-PAOx was analysed using gas chromatography–mass spectrometry. Reaction product peaks are indicated by red arrows. B. Mass spectrum of reaction products and authentic PAOx.

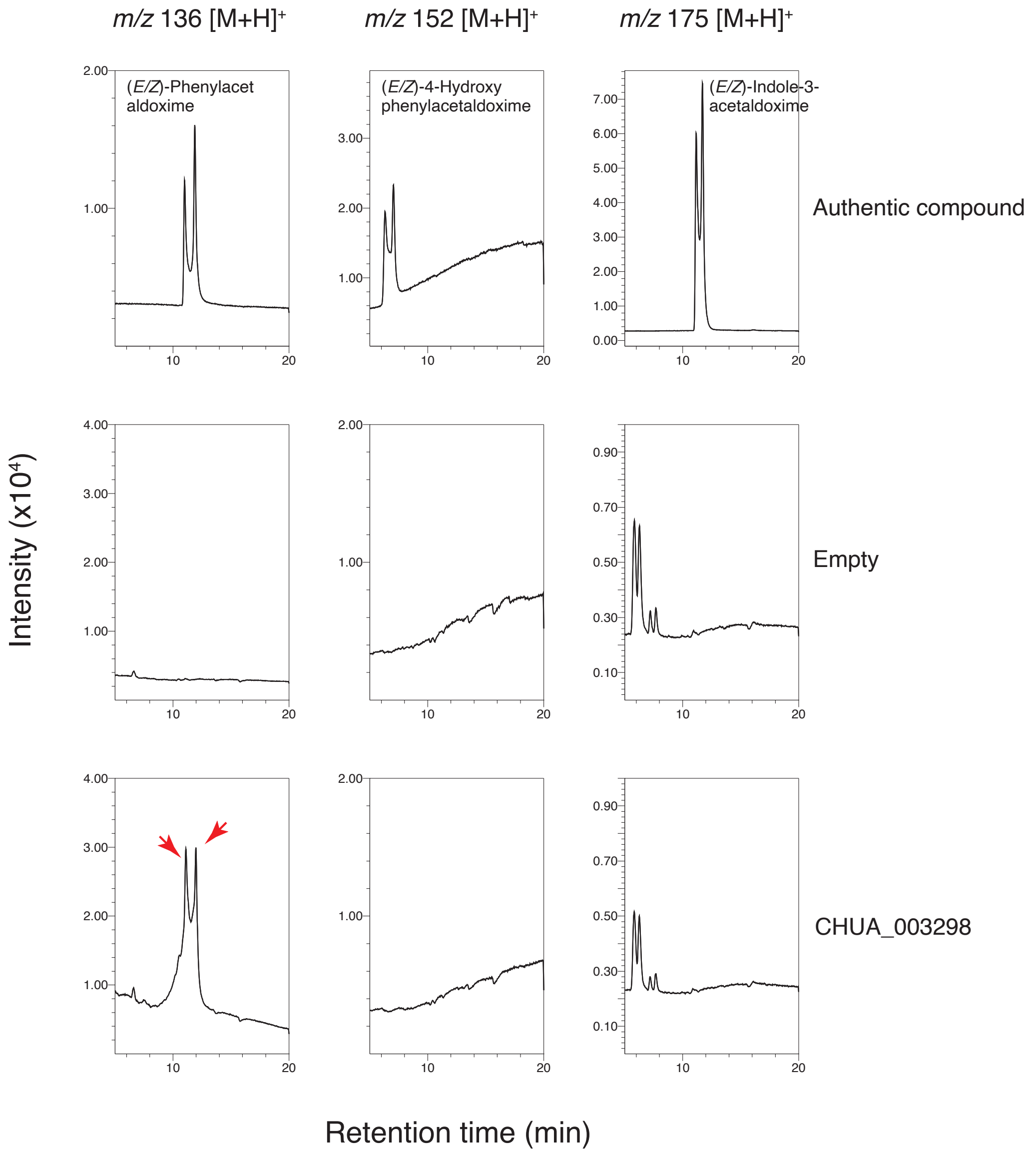

**Fig. S18. Detection of aromatic amino acids-derived aldoximes after the culture of *Escherichia coli* harboring millipede aldoxime synthase (ChuaMOxS, CHUA_003298).** Accumulation of (*E/Z*)-phenylacetaldoxime, (*E/Z*)-4-hydroxyphenylacetaldoxime, and (*E/Z*)-indole-3-acetaldoxime in the medium after the culture of *E. coli* BL21(DE3) carrying pGro7 and empty pET28 plasmid or pET28 carrying CHUA_003298. Selected ion monitoring was used to detect (*E/Z*)-phenylacetaldoxime with *m/z* 136 $[M+H]^+$, (*E/Z*)-4-hydroxyphenylacetaldoxime with *m/z* 152 $[M+H]^+$, and (*E/Z*)-indole-3-acetaldoxime with *m/z* 175 $[M+H]^+$. Aldoximes detected were indicated by red arrows.

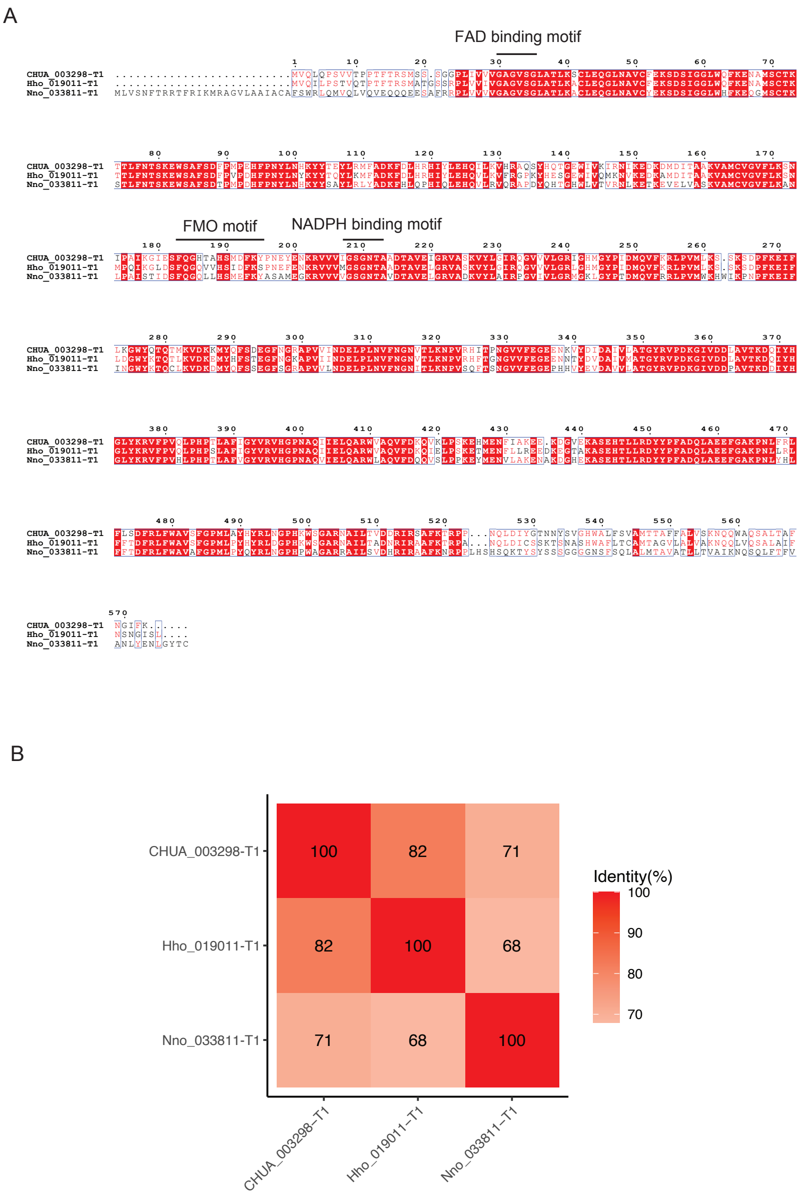

**Fig. S19. Amino acid sequence alignment and amino acid sequence identity of CHUA_003298 (ChuaMOxS) from *Chamberlinius hualienensis*, Hho_019011 from *Helicorthomorpha holstii*, and Nno_033811 from *Niponia nodulosa*.**

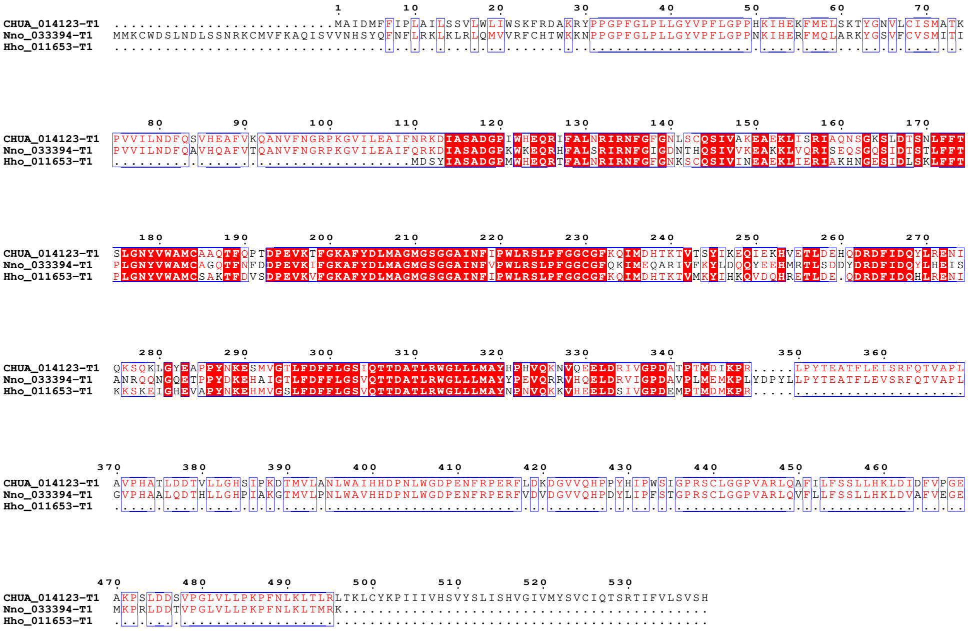

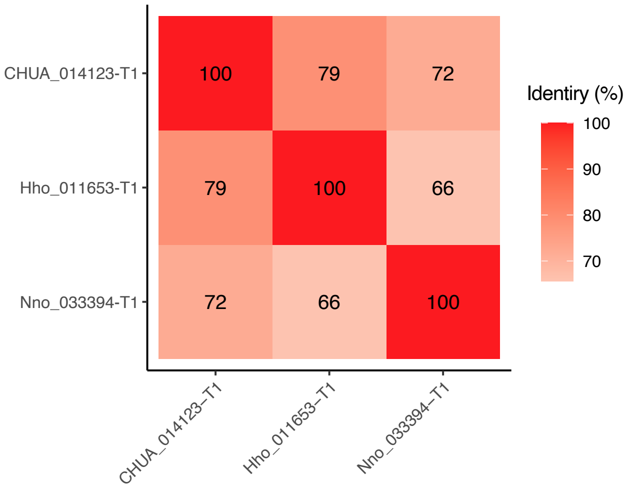

**Fig. S20. Amino acid sequence alignment of CHUA_014123 (CYP30008A2) from *Chamberlinius hualienensis* and its orthorogous proteins, Hho_011653 from *Helicorthomorpha holstii* and Nno_033394 from *Niponia nodulosa*.**

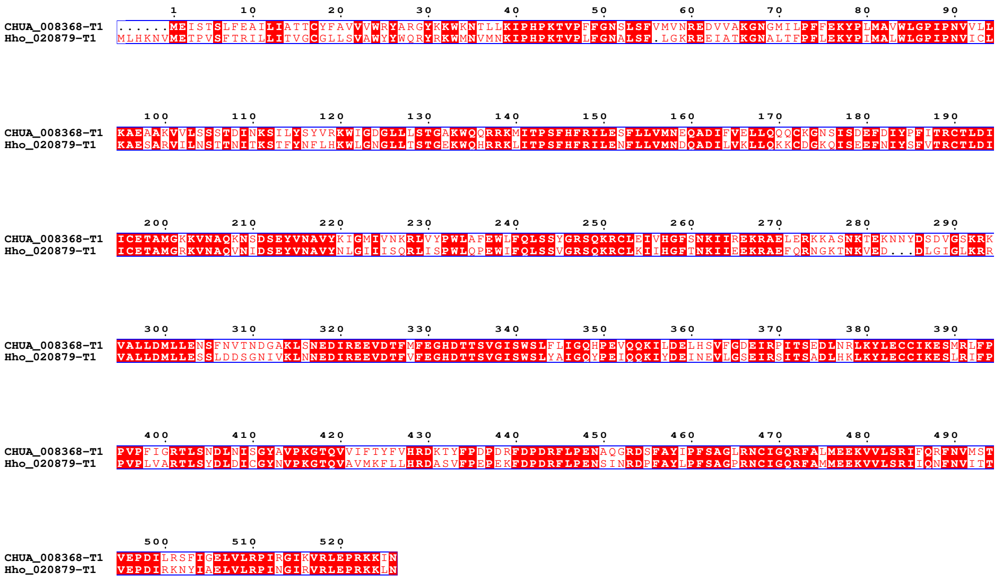

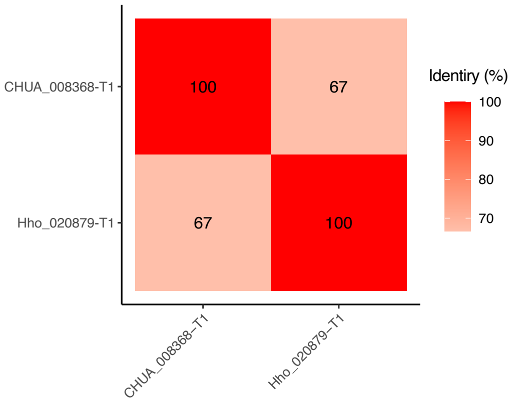

**Fig. S21. Amino acid sequence alignment and amino acid sequence identity of CHUA_008368 (CYP4GL4) from *Chamberlinius hualienensis*, Hho_020879 from *Helicorthomorpha holstii*.**

**Table S1. Shotgun sequencing summary statistics (Illumina)**

| Insert Size | Reads Length | Total Data (Gb) | Sequence coverage |
|---|---|---|---|
| 270 bp | 150:150 | 24.10 | 167 |
| 500 bp | 125:125 | 17.94 | 124 |
| 800 bp | 125:125 | 8.51 | 59 |
| 2 kb | 150:150 | 13.04 | 90 |
| 5 kb | 150:150 | 6.73 | 47 |
| 10 kb | 150:150 | 9.31 | 64 |
| 20 kb | 150:150 | 13.59 | 94 |
| 40 kb | 150:150 | 7.19 | 50 |
| Total | | 100.42 | 695 |

**Table S2. Shotgun sequencing summary statistics (PacBio)**

| Cell | Total Data (Gb) | Average length | Longest read | Shortest length | Sequence coverage |
|---|---|---|---|---|---|
| 1 | 6.94 | 7,078 | 89,590 | 50 | 48 |
| 2 | 8.46 | 9,726 | 73,365 | 50 | 59 |
| 3 | 4.64 | 4,665 | 137,435 | 50 | 32 |
| Total | 20.04 | | | | 139 |

Table S3 TE contents of millipedes and centipedes

Millipede

| Repeat Class/Family | *Chamberlinius hualienensis* | | | *Helicorthomorpha holstii* | | | *Niponia nodulosa* | | | *Anaulaciulus tonginus* | | | *Trigoniulus corallinus* | | | *Glomeris maerens* | | |
|---|---|---|---|---|---|---|---|---|---|---|---|---|---|---|---|---|---|---|
| | Number of elements | Length Occupied (bp) | Percenntage of Sequence (%) | Number of elements | Length Occupied (bp) | Percenntage of Sequence (%) | Number of elements | Length Occupied (bp) | Percenntage of Sequence (%) | Number of elements | Length Occupied (bp) | Percenntage of Sequence (%) | Number of elements | Length Occupied (bp) | Percenntage of Sequence (%) | Number of elements | Length Occupied (bp) | Percenntage of Sequence (%) |
| SINEs | 66 | 4,313 | 0 | 302 | 39,329 | 0.02 | 0 | 0 | 0.00 | 49,684 | 8,661,111 | 1.41 | 141,412 | 22,878,564 | 5.1 | 25 | 2,008 | 0.00 |
| LINEs | 2,115 | 354,534 | 0.25 | 620 | 358,546 | 0.2 | 6,854 | 2,943,860 | 0.90 | 141,032 | 49,660,332 | 8.11 | 194,661 | 73,753,688 | 16.44 | 2,367 | 816,141 | 0.54 |
| LTR elements | 6,262 | 2,226,180 | 1.55 | 7,013 | 6,657,444 | 3.67 | 7,349 | 7,192,089 | 2.19 | 28,489 | 15,014,504 | 2.45 | 15,426 | 7,627,147 | 1.7 | 3,780 | 2,716,145 | 1.81 |
| DNA elements | 24,570 | 4,685,882 | 3.26 | 13,404 | 7,018,807 | 3.87 | 100,750 | 25,771,750 | 7.85 | 60,143 | 16,225,544 | 2.65 | 198,123 | 44,983,912 | 10.03 | 24,248 | 6,197,246 | 4.13 |
| Unclassified | 80,471 | 15,770,108 | 10.99 | 63,394 | 20,346,404 | 11.23 | 321,743 | 72,527,037 | 22.09 | 1,077,794 | 220,597,258 | 36.01 | 266,144 | 61,039,089 | 13.61 | 113,642 | 22,030,347 | 14.70 |
| Total interspersed repeats | 113,484 | 23,041,017 | 16.05 | 84,733 | 34,420,530 | 18.99 | 436,696 | 108,434,736 | 33.03 | 1,357,142 | 310,158,749 | 50.63 | 815,766 | 210,282,400 | 46.88 | 144,062 | 31,761,887 | 14.70 |
| Small RNA | 566 | 45,846 | 0.03 | 419 | 64,421 | 0.04 | 308 | 71,547 | 0.02 | 12,420 | 2,842,939 | 0.46 | 184,120 | 27,607,121 | 6.15 | 540 | 66,017 | 0.04 |
| Satelites | 470 | 85,070 | 0.06 | 24 | 4,793 | 0 | 0 | 0 | 0.00 | 1,057 | 121,424 | 0.02 | 33,767 | 6,632,510 | 1.48 | 246 | 35,256 | 0.02 |
| Simple repeats | 46,168 | 1,765,989 | 1.23 | 1,344 | 471,500 | 0.26 | 71,259 | 2,703,239 | 0.82 | 53,055 | 2,179,604 | 0.36 | 2,407 | 386,405 | 0.09 | 22,079 | 914,042 | 0.61 |
| Low complexity | 10,687 | 501,381 | 0.35 | 247 | 52,179 | 0.03 | 17,046 | 812,313 | 0.25 | 8,300 | 383,178 | 0.06 | 0 | 0 | 0 | 3,192 | 156,608 | 0.10 |
| Total Repeats | 171,375 | 25,439,303 | 17.71 | 86,520 | 34,961,244 | 19.29 | 525,309 | 111,197,075 | 33.87 | 1,431,974 | 318,027,899 | 51.92 | 1,036,060 | 244,908,436 | 54.60 | 170,119 | 33,102,607 | 22.08 |

Centipede

| Repeat Class/Family | *Strigamia maritima* | | | *Rhysida immarginata* | | | *Lithobius niger* | | | *Thereuonema tuberculata* | | |
|---|---|---|---|---|---|---|---|---|---|---|---|---|
| | Number of elements | Length Occupied (bp) | Percenntage of Sequence (%) | Number of elements | Length Occupied (bp) | Percenntage of Sequence (%) | Number of elements | Length Occupied (bp) | Percenntage of Sequence (%) | Number of elements | Length Occupied (bp) | Percenntage of Sequence (%) |
| SINEs | 1,782 | 417,703 | 0.24 | 0 | 0 | 0.00 | 29,126 | 3,438,077 | 0.11 | 0 | 0 | 0.00 |
| LINEs | 3,757 | 1,090,984 | 0.62 | 382,101 | 227,009,366 | 8.98 | 549,785 | 226,602,510 | 7.00 | 349,079 | 155,681,857 | 6.33 |
| LTR elements | 26,920 | 19,739,935 | 11.2 | 69,964 | 54,028,304 | 2.14 | 254,251 | 193,725,228 | 5.98 | 34,993 | 32,513,759 | 1.32 |
| DNA elements | 10,385 | 2,522,989 | 1.43 | 1,332,205 | 442,406,659 | 17.49 | 311,013 | 132,746,291 | 4.10 | 671,250 | 249,445,351 | 10.15 |
| Unclassified | 144,217 | 45,725,301 | 25.95 | 3,547,173 | 783,062,349 | 30.96 | 6,494,229 | 1,415,146,160 | 43.71 | 3,087,097 | 606,764,988 | 24.68 |
| Total interspersed repeats | 187,061 | 69,496,912 | 39.44 | 5,331,443 | 1,506,506,678 | 59.57 | 7,638,404 | 1,971,658,266 | 60.90 | 4,142,419 | 1,044,405,955 | 42.48 |
| Small RNA | 541 | 88,024 | 0.05 | 12,104 | 3,501,496 | 0.14 | 50,168 | 7,576,420 | 0.23 | 56,582 | 19,527,754 | 0.79 |
| Satelites | 1,184 | 378,733 | 0.21 | 21,112 | 5,072,465 | 0.20 | 22,661 | 4,007,979 | 0.12 | 0 | 0 | 0.00 |
| Simple repeats | 1,092 | 232,367 | 0.13 | 394,103 | 15,741,017 | 0.62 | 417,914 | 16,786,819 | 0.52 | 835,139 | 32,018,653 | 1.30 |
| Low complexity | 0 | 0 | 0 | 73,231 | 3,501,085 | 0.14 | 61,755 | 2,930,362 | 0.09 | 183,237 | 8,636,713 | 0.35 |
| Total Repeats | 189,878 | 70,196,036 | 39.83 | 5,831,993 | 1,554,068,493 | 61.45 | 8,190,902 | 2,000,932,573 | 61.80 | 5,217,377 | 1,121,141,731 | 45.61 |

**Table S4. Purification of N-terminal His-tagged ChuaHNL (CHUA_005138) from 1.5 L of *Pichia pastoris* culture supernatant**

| Step | Activity* (U) | Protein (mg) | Specific activity (U/mg) | Yield (%) | Purification fold |
|---|---|---|---|---|---|
| Culture supernatant | 2296 | 636 | 3.6 | 100 | 1 |
| Ni Sepharose | 569 | 0.89 | 643 | 46.8 | 178 |
| MonoQ | 285 | 0.41 | 691 | 23.4 | 191 |

* (*R,S*)-Mandelonitrile degradation activity. Activity was measured with 5 mM (*R,S*)-mandelonitrile.

**Table S5. Purification of N-terminal His-tagged CHUA_005137 from 1.5 L of *Pichia pastoris* culture supernatant**

| Step | Activity* (U) | Protein (mg) | Specific activity (U/mg) | Yield (%) | Purification fold |
|---|---|---|---|---|---|
| Culture supernatant | 258 | 1204 | 0.2 | 100 | 1 |
| Ni Sepharose | 235 | 0.74 | 318 | 19.3 | 1484 |
| MonoQ | 164 | 0.70 | 235 | 13.5 | 1099 |

* (*R,S*)-Mandelonitrile degradation activity. Activity was measured with 5 mM (*R,S*)-mandelonitrile.

**Table S6. Purification of N-terminal His-tagged CHUA_005136 from 1.5 L of *Pichia pastoris* culture supernatant**

| Step | Activity* (U) | Protein (mg) | Specific activity (U/mg) | Yield (%) | Purification fold |
|---|---|---|---|---|---|
| Culture supernatant | 441 | 1152 | 0.38 | 100 | 1 |
| Ni Sepharose | 145 | 4.7 | 30.9 | 11.9 | 81 |
| MonoQ | 81.5 | 1.8 | 45.1 | 6.7 | 118 |

* (*R,S*)-Mandelonitrile degradation activity. Activity was measured with 5 mM (*R,S*)-mandelonitrile.

**Table S7. Characteristics of flavin-dependent monooxygenases (FMOs) from *Chamberlinius hualienensis***

| FMO | Clade | Length | Transmembrane helix* | Localization† |
|---|---|---|---|---|
| CHUA_000006-T1 | Clade II | 558 | 527-546 | Endoplasmic reticulum |
| CHUA_001773-T1 | Clade II | 593 | 571-592 | Endoplasmic reticulum |
| CHUA_003298-T1 (ChuaMOxS) | Clade II | 573 | 511-530 | Endoplasmic reticulum |
| CHUA_008545-T1 | Clade II | 547 | 511-528 | Endoplasmic reticulum |
| CHUA_008546-T1 | Clade II | 539 | 504-522 | Endoplasmic reticulum |
| CHUA_008573-T1 | Clade II | 551 | 528-549 | Endoplasmic reticulum |
| CHUA_010566-T1 | Clade II | 574 | 551-573 | Endoplasmic reticulum |
| CHUA_014222-T1 | Clade II | 542 | 513-531 | Endoplasmic reticulum |
| CHUA_014262-T2 | Clade II | 539 | 237-256, 513-532 | Endoplasmic reticulum |
| CHUA_011563-T1 | Clade I | 423 | | Cytoplasm |

*Predicted using Phobius, † Predicted using Deep Loc

**Table S8 Oligonucleotide primers used in this study**

| Primer | Sequence (5’ – 3’) |
|---|---|
| For cloning and construction of ChuaCPR expression plasmid | |
| GbSc-ChuaCPR-FW | cacactaaattaccggatccaaaaaaATGGCCGCCGAAGAAGTTTTGGATTC |
| GbSc-ChuaCPR-RV | atcccccgcggaattcTTAACTCCAGACATCAGCTGAATATC |
| Gb-GALP-FW | tgttgaagcttgcatgcGATCAAAAATCATCGCTTCGC |
| Gb-PGKT-RV | tcgacctgcaggcatgcTTCGAAACGCAGAATTTTCG |
| For cloning and construction of ChuaCYP expression plasmid | |
| GbSc-3201B1-FW | cacactaaattaccggatccaaaaaaATGGCGCCGAGTGTAGATCAGTGG |
| GbSc-3201B1-RV | atcccccgcggaattcTTAATCTTTTCTAATTTGAATTAAAAG |
| GbSc-8368-FW | cacactaaattaccggatccaaaaaaATGGAGATTTCTACAAGCTTATTTG |
| GbSc-8368-RV | atcccccgcggaattcTTAATTTATTTTTTTTCTTGGCTCC |
| GbSc-10648-FW | cacactaaattaccggatccaaaaaaATGGATCTAAGTTACGTTTGGTCG |
| GbSc-10648-RV | atcccccgcggaattcTCAATCTTCTCGTAATTTGATTAG |
| GbSc-3521-FW | cacactaaattaccggatccaaaaaaATGAAATTAGACGATATTAATTAC |
| GbSc-3521-RV | atcccccgcggaattcTCTAACATGAACTACCAACGAATA |
| GbSc-3525-FW | cacactaaattaccggatccaaaaaaATGATTTTTGAAGCGTACGCAGGA |
| GbSc-3525-RV | atcccccgcggaattcTTATTCGTTTCTACTGTTTATAAG |
| GbSc-14123-FW | cacactaaattaccggatccaaaaaaATGGCCATAGACATGTTTTTTATC |
| GbSc-14123-RV | atcccccgcggaattcTTATAATCTGAGTGTCAGCTTCAA |
| GbSc-7584-FW | cacactaaattaccggatccaaaaaaATGGAGTTGGGAATAATTTCTTGG |
| GbSc-7584-RV | atcccccgcggaattcTTAGTCAAGCACTCTTTGTTTGGC |
| GbSc-7622-FW | cacactaaattaccggatccaaaaaaATGGTATGTGATCAACTGATGGCA |
| GbSc-7622-RV | atcccccgcggaattcTTAACTTCGAAGTACTGCACAAAT |
| GbSc-10928-FW | cacactaaattaccggatccaaaaaaATGTCTCTAGCGCCCATAATCGCA |
| GbSc-10928-RV | atcccccgcggaattcTCATTCTCTCTGTTCCAGTTTTAC |
| GbSc-10946-FW | cacactaaattaccggatccaaaaaaATGGCGTTCGTCGACGTGTGGACT |
| GbSc-10946-RV | atcccccgcggaattcTTATTCATTTCTAGTTTCAATTAA |
| GbSc-821-FW | cacactaaattaccggatccaaaaaaATGATGACTAATTATGTTCATGGG |
| GbSc-821-RV | atcccccgcggaattcTTATCTTTCGTGAAATTTAAGCTT |
| GbSc-348-FW | cacactaaattaccggatccaaaaaaATGGCTTTAGAAGAACTTTTCACA |
| GbSc-348-RV | atcccccgcggaattcTTAGATTTTCTCCCTCTCTCTGAC |
| GbSc-1815-FW | cacactaaattaccggatccaaaaaaATGCTTTTTGCAACTATTTTTAGT |

| | |
|---|---|
| GbSc-1815-RV | atcccccgcggaattcTTAACATTTTCTTAATATTGCAGA |
| GbSc-1891-FW | cacactaaattaccggatccaaaaaaATGTCTATTGCTCTATTACTATTA |
| GbSc-1891-RV | atcccccgcggaattcTTATAAATCTTCAAATATAACCCA |
| GbSc-1921-FW | cacactaaattaccggatccaaaaaaATGGATAACCTTGGCTTTATTATT |
| GbSc-1921-RV | atcccccgcggaattcCTAATTTCGTCTCTTGACATATAA |
| GbSc-11710-FW | cacactaaattaccggatccaaaaaaATGGATATTTTCGGATTTTTTTTG |
| GbSc-18710-RV | atcccccgcggaattcTTAACTTCGTTTTTCAACAGTCAA |
| GbSc-8576-FW | cacactaaattaccggatccaaaaaaATGATTGAATTGAACACGATCAGT |
| GbSc-8576-RV | atcccccgcggaattcTTATCTTTGTTGTACCTTGACATG |
| GbSc-8495-T2-FW | cacactaaattaccggatccaaaaaaATGGATTTGTGGACAGCATTTGTC |
| GbSc-8495-T2-RV | atcccccgcggaattcTTAGTGTCTAGATTTGATACAAAT |
| GbSc-8495-T1-FW | cacactaaattaccggatccaaaaaaATGGATTTCTTGAGTATATTAGGA |
| GbSc-8495-T1-RV | atcccccgcggaattcTTATTTCGGTCTTGACTTGACACA |
| GbSc-8467-FW | cacactaaattaccggatccaaaaaaATGTGGTTAATAATTGTGATTTCT |
| GbSc-8467-RV | atcccccgcggaattcTCATCGGATGTTGATCTTTAGCGC |
| GbSc-9816-FW | cacactaaattaccggatccaaaaaaATGTTTGCTGCAATACAACTACTC |
| GbSc-9816-RV | atcccccgcggaattcTTATCGCTTTTTTAATGAGATCCA |
| GbSc-6572-FW | cacactaaattaccggatccaaaaaaATGTGGTTGGAGAGCGCGCTGCTG |
| GbSc-6572-RV | atcccccgcggaattcTTACTTGCATCTGCTTTTCAAAAC |
| GbSc-12471-FW | cacactaaattaccggatccaaaaaaATGGAGTTATTAGGAATAATTGAT |
| GbSc-12471-RV | atcccccgcggaattcTTATTTTCTGGGTTCAATTTTTAC |
| GbSc-3524-FW | cacactaaattaccggatccaaaaaaATGTTCGAAGTGAATGCATGTGAA |
| GbSc-3524-RV | atcccccgcggaattcTTATTCTTTTCTATGGTTAACAAG |
| GbSc-11710-FW | cccgggtaccaaaaaaATGGATATTTTCGGATTTTTTTTGA |
| GbSc-11710-RV | atcccccgcggaattcTTAACTTCGTTTTTCAACAGTCAAA |
| GbSc-727-FW | cccgggtaccaaaaaaATGGTGTTCAATGCCTTACTTCAGC |
| GbSc-727-RV | atcccccgcggaattcCTAATTTTCTCGTTTTTCGAAGCAA |
| GbSc-8368-FW | cccgggtaccaaaaaaATGGAGATTTCTACAAGCTTATTTG |
| GbSc-8368-RV | atcccccgcggaattcTTAATTTATTTTTTTTCTTGGCTCC |
| GbSc-3201B2-FW | cacactaaattaccggatccaaaaaaATGTTGGGTGAATATTTAAAGTTG |
| GbSc-3201B2-RV | atcccccgcggaattcTCATCGCAATTGTACCAAAAGTTT |
| GbSc-4GL3-FW | cacactaaattaccggatccaaaaaaATGGATTTCAACGCCATTCGAGTT |
| GbSc-4GL3-RV | atcccccgcggaattcTTAATCTCTAAGTTCAAGTTGAAC |
| GbSc-4GL2-FW | cacactaaattaccggatccaaaaaaATGTTGTTTATAATTAGCGTTATA |

| | |
|---|---|
| GbSc-4GL2-RV | atcccccgcggaattcTCATAGACCATTTCTCGCTTCTAG |
| GbSc-4GL1-FW | cacactaaattaccggatccaaaaaaATGGCTGTAATTAGTGATTTTGTC |
| GbSc-4GL1-RV | atcccccgcggaattcCTAAATCTTTGGCCTGGGCTGCAA |
| GbSc-3201E1-FW | cacactaaattaccggatccaaaaaaATGAATTTGGAAGCGATCGCCTAC |
| GbSc-3201E1-RV | atcccccgcggaattcCTAATACTTTCTAACTGCGAATGT |
| GbSc-3201G1-FW | cacactaaattaccggatccaaaaaaATGGCGTTTTACATATGGTCGTTT |
| GbSc-3201G1-RV | atcccccgcggaattcTTAATCCTCTCTCAATTCGATTAA |
| GbSc-3201G2-FW | cacactaaattaccggatccaaaaaaATGGAAGTTGGCTACGTGTGGTCG |
| GbSc-3201G2-RV | atcccccgcggaattcTTAATCATTTCTCAATTGAATTAA |
| GbSc-3201G3-FW | cacactaaattaccggatccaaaaaaATGGATCTAAGTTACGTTTGGTCG |
| GbSc-3201G3-RV | atcccccgcggaattcTTAAATTTCTCGTAATTTGATTAG |
| GbSc-3201G4-FW | cacactaaattaccggatccaaaaaaATGGATCTAAGTTACGTTTGGTCG |
| GbSc-3201G4-RV | atcccccgcggaattcTTAAATTTCTCGTAATTTGATTAG |
| GbSc-3201F1-FW | cacactaaattaccggatccaaaaaaATGTCTCTAATTGAAAAAGCAATT |
| GbSc-3201F1-RV | atcccccgcggaattcTTAGCGAGGTTGAAGAAACAGTGA |
| GbSc-3198A1-FW | cacactaaattaccggatccaaaaaaATGTTAAACGAAATTTTAAATTCA |
| GbSc-3198A1-RV | atcccccgcggaattcTTAAAGGTTTCTGCGTTTCATGAT |
| GbSc-4GN1-FW | cacactaaattaccggatccaaaaaaATGACTTCTGTGCTTGTTTTAGTT |
| GbSc-4GN1-RV | atcccccgcggaattcTTAATATTCTTTCTCAATTGGTTT |
| GbSc-4GN2-FW | cacactaaattaccggatccaaaaaaATGGCTTCCATGCTGTTACTTGCT |
| GbSc-4GN2-RV | atcccccgcggaattcTTAGTTTTCCGTCTCGATTGGTTT |
| GbSc-3200B1-FW | cacactaaattaccggatccaaaaaaATGGATGCGTGGACCGTATTAGCC |
| GbSc-3200B1-RV | atcccccgcggaattcTCAGTGTCTTGATTTGATACATAT |
| GbSc-3200B2-FW | cacactaaattaccggatccaaaaaaATGGATGTAGTGAGTTTTGGCACA |
| GbSc-3200B2-RV | atcccccgcggaattcTTATCTTGCCTTGATGCAAATAAA |
| GbSc-3200B3-FW | cacactaaattaccggatccaaaaaaATGGATTTATGGAACATATTTGCATG |
| GbSc-3200B3-RV | atcccccgcggaattcTTATCGTGATTTAATAGAAATTAG |
| GbSc-3200B4-FW | cacactaaattaccggatccaaaaaaATGGATTTGTGGAGCATACTAACA |
| GbSc-3200B4-RV | atcccccgcggaattcTTATCTTGTTTTAATAGAAATTAG |
| GbSc-3200D1-FW | cacactaaattaccggatccaaaaaaATGGTGGCCATCATACTTATTGG |
| GbSc-3200D1-RV | atcccccgcggaattcTCAGTCTTGTCGTGGTACCAAACA |
| GbSc-3201C1-FW | cacactaaattaccggatccaaaaaaATGGCGTGGCTACCAAGCAGTGGG |
| GbSc-3201C1-RV | atcccccgcggaattcTTATGATTTTCTACATTTGACATA |
| GbSc-3201A1-FW | cacactaaattaccggatccaaaaaaATGGTTGCTATGATAAATTCAGAG |

| | |
|---|---|
| GbSc-3201A1-RV | atcccccgcggaattcTTAGTTAACAAATCGTTTAGTAAT |
| GbSc-14000-FW | cacactaaattaccggatccaaaaaaATGTCGCCTGTTTTGTTTGGAATA |
| GbSc-14000-RV | atcccccgcggaattcTCAATGTGTTTTTTCAACTAATTG |
| GbSc-4GQ1-FW | cacactaaattaccggatccaaaaaaATGATCATAAGTGCTTTTGTAGCA |
| GbSc-4GQ1-RV | atcccccgcggaattcTTATCTCTTTTCCAAACTAATGAG |
| GbSc-11710-FW | cacactaaattaccggatccaaaaaaATGGATATTTTCGGATTTTTTTTG |
| GbSc-11710-RV | atcccccgcggaattcTTAACTTCGTTTTTCAACAGTCAA |
| GbSc-2527-FW | cacactaaattaccggatccaaaaaaATGAAGTTCTTATCATCAATTGTC |
| GbSc-2527-RV | atcccccgcggaattcTTAGATTTTGGGGATTAATTGTTT |
| GbSc-8909-FW | cacactaaattaccggatccaaaaaaATGACACCACTTGACGTTTGCTGG |
| GbSc-8909-RV | atcccccgcggaattcTCAACATTGTGGTCTAGTTAGAGC |
| GbSc-9818-FW | cacactaaattaccggatccaaaaaaATGCAAGCTCTCTTGCAACTCTTG |
| GbSc-9818-RV | atcccccgcggaattcTTAATGCCGATTTTCAAGTTTAAC |
| GbSc-3193A1-FW | cacactaaattaccggatccaaaaaaATGGAGTTGTTTGGCTTAATT |
| GbSc-3193A1-RV | atcccccgcggaattcTTAATTTCGTCTTTCAACTTTGAG |
| GbSc-5942-FW | cacactaaattaccggatccaaaaaaATGGAGTTGTTCGGCTTAATC |
| GbSc-5942-RV | atcccccgcggaattcTTAATTTCGTCTTTCGACTTTGAG |
| GbSc-5727-FW | cacactaaattaccggatccaaaaaaATGGATTTGGAGTGGAACCAACTT |
| GbSc-5727-RV | atcccccgcggaattcTTATCGTCGTTCAATTTTTACGTA |
| GbSc-5499-FW | cacactaaattaccggatccaaaaaaATGGATATCAGACATATTTGGGAA |
| GbSc-5499-RV | atcccccgcggaattcCTAATCATTTTGAATTATAGGTAT |
| GbSc-4975-FW | cacactaaattaccggatccaaaaaaATGATTTTTTGGTGGACAGTCGTT |
| GbSc-4975-RV | atcccccgcggaattcTTAAGGTCTTAGAGTGTGATTTTG |
| GbSc-1635-FW | cacactaaattaccggatccaaaaaaATGGCAGCGAGTTACGTTTGGTCG |
| GbSc-1635-RV | atcccccgcggaattcTTAATCATATCTCACTTCAATTAA |
| GbSc-4GM1-FW | cacactaaattaccggatccaaaaaaATGTTAAACATTTTATTTTTCTTT |
| GbSc-4GM1-RV | atcccccgcggaattcCTAATTAGTTGCTCTCGACTTAAA |
| GbSc-4GM2-FW | cacactaaattaccggatccaaaaaaATGATGATAACCGTGTCTTGTTAC |
| GbSc-4GM2-RV | atcccccgcggaattcTTATGTAGTTCGCCTTGGCTGAAT |
| GbSc-18G1-FW | cacactaaattaccggatccaaaaaaATGATGAATACATTGCAGAACATT |
| GbSc-18G1-RV | atcccccgcggaattcTTAAAAATGAAATCTTGGAATTGC |
| GbSc-3200C1-FW | cacactaaattaccggatccaaaaaaATGGCTGTAGAGGAGCTTTTCAGA |
| GbSc-3200C1-RV | atcccccgcggaattcTTAGATTTTCTCCCTCTCTCTAAC |
| GbSc-3200C3-FW | cacactaaattaccggatccaaaaaaATGGCGGTTGATTTCATCTGGTTA |

| | |
|---|---|
| GbSc-3200C3-RV | atcccccgcggaattcCTATTGCCGTAATATTGCATGGAC |
| GbSc-3200C4-FW | cacactaaattaccggatccaaaaaaATGGCGATTGATCAAAATTTAATG |
| GbSc-3200C4-RV | atcccccgcggaattcTCATCGAGCAACTGCGCAAATATT |
| GbSc-3200C5-FW | cacactaaattaccggatccaaaaaaATGGGAGTTGATTCCAATATAACA |
| GbSc-3200C5-RV | atcccccgcggaattcTTAACATCGAGAAATTGCACAAAC |
| GbSc-3200C6-FW | cacactaaattaccggatccaaaaaaATGTGGACATTCTATTTGGCAATT |
| GbSc-3200C6-RV | atcccccgcggaattcCTACTTTCGTGGTTTTGCACAAAC |
| GbSc-7618-FW | cacactaaattaccggatccaaaaaaATGGCAGCTGATTACAATTTAATC |
| GbSc-7618-RV | atcccccgcggaattcTTATCGTCGAGATATTGCACAAAT |
| GbSc-3200A1-FW | cacactaaattaccggatccaaaaaaATGGCTGTAGAGGAGCTTTTCAGA |
| GbSc-3200A1-RV | atcccccgcggaattcTTAGATTTTCTCCCTCTCTCTAAC |
| GbSc-3200A2-FW | cacactaaattaccggatccaaaaaaATGGCTTTAGAAGAGCTTTTTATC |
| GbSc-3200A2-RV | atcccccgcggaattcCTATTCGGTAGACCTCTCTCTGAC |
| GbSc-1922-FW | cacactaaattaccggatccaaaaaaATGACTGTTGTTACTACTTCCGAT |
| GbSc-1922-RV | atcccccgcggaattcTTATCGTTTAGTAATGTACAATGA |
| GbSc-4GP1-FW | cacactaaattaccggatccaaaaaaATGGCTAGTTGGTTAATTTATGCA |
| GbSc-4GP1-RV | atcccccgcggaattcTTAATGCGATTTCCTATTCTCCAA |
| GbSc-8345-FW | cacactaaattaccggatccaaaaaaATGTTAAGTCTGAGTCACTTTATT |
| GbSc-8345-RV | atcccccgcggaattcTTAAAGCTCTTTGCCAGTTGTTTT |
| GbSc-9817-FW | cacactaaattaccggatccaaaaaaATGCAGACGGTAACTGTATTTCTA |
| GbSc-9817-RV | atcccccgcggaattcTTACAAAAAAAACCGGTCAAAGTT |
| GbSc-10635-FW | cacactaaattaccggatccaaaaaaATGAATCTAAGTTACGTTTGGTCG |
| GbSc-10635-RV | atcccccgcggaattcTTAATTTTCTCGTGATTTGATTAG |
| GbSc-13654-FW | cacactaaattaccggatccaaaaaaATGTCATTGAAGCTGTATAAAGCC |
| GbSc-13654-RV | atcccccgcggaattcTTAATCTTCTCTCAATTCGATTAA |
| GbSc-306B1-FW | cacactaaattaccggatccaaaaaaATGTTGCCGCTGAGATTTCAG |
| GbSc-306B1-RV | atcccccgcggaattcTTATCTCAAAACCGCTCGCAACAT |
| GbSc-3546-FW | cacactaaattaccggatccaaaaaaATGCCTAACATGCTCGGGCTA |
| GbSc-3546-RV | atcccccgcggaattcTTAATCGAAATTGATAAAATTCGG |
| GbSc-3547-FW | cacactaaattaccggatccaaaaaaATGATATTCGAATTATTCACA |
| GbSc-3547-RV | atcccccgcggaattcTCAAAGGCTTCCATTTTTATTTGC |
| GbSc-3200C1-FW | cacactaaattaccggatccaaaaaaATGGAGGTATCTACCAATCCT |
| GbSc-3200C1-RV | atcccccgcggaattcTCAGAATGGATATTCACGTAAAAC |
| GbSc-3201D1-FW | cacactaaattaccggatccaaaaaaATGGAACTGTGGTGGTTTTGG |

| | |
|---|---|
| GbSc-3201D1-RV | atcccccgcggaattcTTAGTTTCTAATTTTAACGAGTAA |
| GbSc-4GP1-FW | cacactaaattaccggatccaaaaaaATGGCTAGTTGGTTAATTTAT |
| GbSc-4GP1-RV | atcccccgcggaattcTTAATGCGATTTCCTATTCTCCAA |
| GbSc-6737-FW | cacactaaattaccggatccaaaaaaATGAATGTTTTAGATGGTTTT |
| GbSc-6737-RV | atcccccgcggaattcCTATTTTCTTTTCTGGATACCAAC |
| GbSc-13084-FW | cacactaaattaccggatccaaaaaaATGTTGGAAATGCCGTCACTT |
| GbSc-13084-RV | atcccccgcggaattcTTATATTGCTCGTTTAACGAG |
| GbSc-10648-FW | cacactaaattaccggatccaaaaaaATGGATCTAAGTTACGTTTGG |
| GbSc-10648-RV | atcccccgcggaattcTTAAATTTCTCGTAATTTGAT |
| GbSc-2961-FW | cacactaaattaccggatccaaaaaaATGAATTTGGAAGCGATCGCC |
| GbSc-2961-RV | atcccccgcggaattcCTAATACTTTCTAACTGCGAA |
| GbSc-8097-FW | cacactaaattaccggatccaaaaaaATGGCGTTTTACATATGGTCG |
| GbSc-8097-RV | atcccccgcggaattcTTAATCCTCTCTCAATTCGAT |
| GbSc-6739-FW | cacactaaattaccggatccaaaaaaATGAATGTATTGGGTTTATTA |
| GbSc-6739-RV | atcccccgcggaattcTTATGTAGATCTTGTTTCAAC |
| GbSc-6738_T2-FW | cacactaaattaccggatccaaaaaaATGGATGTTTTTGGATTGGTT |
| GbSc-6738_T2-RV | atcccccgcggaattcCTATTTCCTTGGTTGAATTTT |
| For cloning of FMO | |
| ChuaFMO6-FW | ATGGATATCGACAGTGAATCTAAA |
| ChuaFMO6-RV | TCATTGATGTATTGTCATTAGACC |
| ChuaFMO1773-FW | ATGTCTTACGAAAGATCTAAGAAG |
| ChuaFMO1773-RV | TTAAAACAAATAGAAACATGAGAA |
| ChuaFMO3298-FW | ATGGTTCAACTACAGCCATCCGTC |
| ChuaFMO3298-RV | TTATTTAAAGATGCCATTGAAAGCTGT |
| ChuaFMO8545-FW | ATGGCATACACTGTAGATGTTGTG |
| ChuaFMO8545-RV | TCAGCGATTTTTGACTGCCATTAC |
| ChuaFMO8546-FW | ATGGCTGTTGCAGTAGATGTTGTG |
| ChuaFMO8546-RV | TTACCTAACATCATGCAGTAGTAA |
| ChuaFMO8573-FW | ATGAAGCCTGTGGTGATTATTGGA |
| ChuaFMO8573-RV | TTAGTTAGCTATAAAAACGTAAAT |
| ChuaFMO10566-FW | ATGAAATCAATGGACATTTCGAGT |
| ChuaFMO10566-RV | TTATAAGACAAACATATAAGCAAT |
| ChuaFMO14222-FW | ATGGCGGCGATTGAGGTGGACGTG |

| | |
|---|---|
| ChuaFMO14222-RV | TTATTTAGCAAATATTTCATTCAA |
| ChuaFMO14262-FW | ATGACGACGTTTGATGTAGTCGTA |
| ChuaFMO14262-RV | TCAAAAGCGCGCCGACAAATCGCG |
| ChuaFMO-11563-FW | ATGAAGGTTTGTATAATTGGAGCTGGT |
| ChuaFMO-11563-RV | TCATCGATAGAAGACACAATTGTCTCT |
| For construction of *Escherichia coli* expression plasmids | |
| GbEc-ChuaFMO6-FW | TGCCGCGCGGCAGCCAtATGGATATCGACAGTGAATCTAAA |
| GbEc-ChuaFMO6-RV | tcgagtgcggccgcaagcttaTCCCGAAGATTTTTTTTCTATTTT |
| GbEc-ChuaFMO1773-FW | TGCCGCGCGGCAGCCAtATGTCTTACGAAAGATCTAAGAAG |
| GbEc-ChuaFMO1773-RV | tcgagtgcggccgcaagcttaATTAGCGATCAGTTTACTTAATTC |
| GbEc-ChuaFMO3298-FW | TGCCGCGCGGCAGCCAtATGGTTCAACTACAGCCATCCGTC |
| GbEc-ChuaFMO3298-RV | TCGAGTGCGGCCGCaagcttTTATTTAAAGATGCCATTGAAAGC |
| GbEc-ChuaFMO8545-FW | TGCCGCGCGGCAGCCAtATGGCATACACTGTAGATGTTGTG |
| GbEc-ChuaFMO8545-RV | tcgagtgcggccgcaagcttaGCATTTCCCAGTTTTTTGCTTACA |
| GbEc-ChuaFMO8546-FW | TGCCGCGCGGCAGCCAtATGGCTGTTGCAGTAGATGTTGTG |
| GbEc-ChuaFMO8546-RV | tcgagtgcggccgcaagcttaCTTTCTAATACGTTGTTGTTTACG |
| GbEc-ChuaFMO8573-FW | TGCCGCGCGGCAGCCAtATGAAGCCTGTGGTGATTATTGGA |
| GbEc-ChuaFMO8573-RV | tcgagtgcggccgcaagcttaTGGATTGTTGTAGTTGGTCTCCAT |
| GbEc-ChuaFMO10566-FW | TGCCGCGCGGCAGCCAtATGAAATCAATGGACATTTCGAGT |
| GbEc-ChuaFMO10566-RV | tcgagtgcggccgcaagcttaAGAAGACTGTTTACAGCAATTGTT |
| GbEc-ChuaFMO14222-FW | TGCCGCGCGGCAGCCAtATGGCGGCGATTGAGGTGGACGTG |
| GbEc-ChuaFMO14222-RV | tcgagtgcggccgcaagcttaTCTCTTCTTTTTACGCCCATTATT |
| GbEc-ChuaFMO14262-FW | TGCCGCGCGGCAGCCAtATGACGACGTTTGATGTAGTCGTA |
| GbEc-ChuaFMO14262-RV | tcgagtgcggccgcaagcttaGCGAGTATGACTGTTTTTATCATC |
| GbEc-ChuaFMO11563-FW | cacactaaattaccggatccaaaaaaATGAAGGTTTGTATAATTGGAGCT |
| GbEc-ChuaFMO11563-FW | atcccccgcggaattcTCATCGATAGAAGACACAATTGTC |

**Table S9. RNA-seq for gene annotation**

| Tissue | Clean Reads (million) | Clean bases (Gbp) | Reads Length | Q30(%) | GC (%) |
|---|---|---|---|---|---|
| Antennae (male) | 44.1 | 6.6 | 150:150 | 93 | 39 |
| Antennae (female) | 42.3 | 6.3 | 150:150 | 93 | 38 |
| Segments with defensive glands | 50.6 | 7.6 | 150:150 | 95 | 38 |
| Segments without defensive glands | 49.7 | 7.4 | 150:150 | 95 | 39 |
| Gut | 44.8 | 6.7 | 150:150 | 94 | 39 |

**Table S10. RNA-seq for gene expression analysis**

| Tissue | Clean Reads (million) | Clean bases (Gbp) | Reads Length | Q30 (%) | GC (%) |
|---|---|---|---|---|---|
| Antennae (male)-1 | 11.3 | 3.4 | 150:150 | 91.83 | 39 |
| Antennae (male)-2 | 10.8 | 3.2 | 150:150 | 91.95 | 39 |
| Antennae (male)-3 | 9.7 | 2.9 | 150:150 | 91.16 | 39 |
| Paraterga with defensive glands-1 | 11.9 | 3.6 | 150:150 | 91.28 | 39 |
| Paraterga with defensive glands-2 | 11.4 | 3.4 | 150:150 | 92.34 | 39 |
| Paraterga with defensive glands-3 | 11.4 | 3.4 | 150:150 | 92.25 | 40 |
| Paraterga without defensive glands-1 | 11.6 | 3.5 | 150:150 | 91.68 | 39 |
| Paraterga without defensive glands-2 | 12.3 | 3.7 | 150:150 | 91.86 | 39 |
| Paraterga without defensive glands-3 | 11.1 | 3.3 | 150:150 | 92.38 | 40 |
| Gut-1 | 10.8 | 3.2 | 150:150 | 92.10 | 39 |
| Gut-2 | 12.1 | 3.6 | 150:150 | 92.04 | 40 |
| Gut-3 | 10.4 | 3.1 | 150:150 | 91.55 | 39 |
